# Nrl expression and promoter activity in developing cone photoreceptors

**DOI:** 10.64898/2026.08.06.743042

**Authors:** Brandon P. Webley, Angelina V. Grebe, Brandon Mohammed, Jhon Lopez, Afif Zaman, Miruna Ghinia-Tegla, Mark M. Emerson

## Abstract

The MAF family transcription factor *Nrl* is a key regulator of vertebrate rod photoreceptor formation/differentiation. The expression of *Nrl* in nascent photoreceptors has been proposed to act as a master regulator to initiate a rod program and to repress a default cone photoreceptor sister fate. The evidence for rod-specific expression of *Nrl* has been most strongly supported by mouse transgenic models that use a transcriptional promoter element for *Nrl*. However, this element has not been rigorously evaluated to have rod lineage-specific activity or as a validated proxy for endogenous *Nrl* expression, despite its wide use under this assumption. Here we identify that the *Nrl* promoter is not exclusively active in rod photoreceptors but is also transiently active within a high proportion of the differentiating cone photoreceptor population. Thus, the *Nrl* promoter cannot be used to distinguish rods from cones at developmental timepoints. Furthermore, multiple lines of evidence using immunofluorescence and reanalysis of published single cell RNA-seq/ATAC-seq datasets support the presence of endogenous *Nrl* gene expression in early differentiating cone photoreceptors. Taken together, these results suggest that proposed gene regulatory networks that position *Nrl* as a master regulator of rod versus cone fate are inadequate as currently constructed. This impacts our understanding of visual system evolution and has implications for the development of therapeutic strategies of cell replacement for human blindness.

## Introduction

The majority of vertebrate retinas contain two photoreceptor types that mediate conscious vision – rods and cones^1^. Cones are integral for high acuity visual information and mediate color vision, while rods are sensitive to very low levels of light, mediating scotopic vision^2^. Together, these two photoreceptor types enable visual capabilities across a broad range of light conditions, and cone loss through autonomous or non-cell autonomous mechanisms leads to a dramatic reduction in visual capacity^3^.

When photoreceptors degenerate in humans, such as in disorders like retinitis pigmentosa and age-related macular degeneration, there is no way to replenish them^4^. Affected individuals may eventually totally lose their sight and this can lead to significant diminishment of quality of life as well as have a severe impact on an individual’s life expectancy. Global estimates predict that by 2050 approximately 474 million people will be living with moderate or severe vision impairment, underscoring the growing burden of ocular disease worldwide ^5^.

The *Nrl* gene encodes a large MAF family transcription factor, which has been identified in human patients as a genetic contributor to rod degeneration in retinitis pigmentosa, also leading to irreversible loss of cones and eventual blindness ^6,7^. *Nrl* has been reported in previous studies to be specifically expressed in differentiating and mature rod photoreceptors and is the only large MAF protein expressed in the mouse and human neural retina^8–10^. In mice, loss of *Nrl* has also been shown to cause similar phenotypes as those found in humans ^11,12^. A previous report found that *Nrl* expression driven from a *Crx* promoter in both WT and *Nrl* KO contexts leads to loss of *Arr3* expression in mouse cones, and this included loss of other cone phototransduction genes as determined by RT-PCR^13^. Expression of *Nrl* driven from the *Opn1sw* promoter, BPp, in the *Nrl* KO background, induced rhodopsin in S-opsin expressing cells. *Nrl* has since been described as a “master regulator” of rod fate commitment^14–18^. The primacy of *Nrl* in rod photoreceptor differentiation has made understanding the genetic mechanisms that control *Nrl* expression crucial for elucidating the cause and progression of photoreceptor degeneration in retinitis pigmentosa.

Transcriptional regulation has been the primary mechanistic model by which *Nrl* expression induces the rod photoreceptor fate. A seminal study reported that a 2.5kb fragment of the *Nrl* promoter was capable of recapitulating NRL expression and was consistent with the transcriptional regulation of *Nrl* being limited to rod photoreceptors^8^. The same report noted that fate-mapping experiments using the *Nrl* promoter driving *Cre* supported the finding that the promoter is rod-specific (listed as unpublished data). A subsequent study remade the 2.5kb *Nrl* promoter driving *Cre* transgenic mouse line, which will be referred to as *NrlpCre* ^19^. The *NrlpCre* line was crossed to a *Cre*-responsive reporter line and the study reported that the lineage traced cells did not localize with the cone marker cone-arrestin but labeled the majority of the ONL, leading to a conclusion that the *NrlpCre* drives *Cre*-recombination specifically in developing rods. However, this study did not employ quantitative methods to characterize the activity of this element. Taken together, the use of the *Nrl* promoter as a proxy for *NRL* expression and/or of rod photoreceptor cell identity through developmental time has not been rigorously demonstrated in these studies.

In this report, evidence is presented that challenges the findings in the original *NrlpGFP* report, Akimoto et al 2006, as well as subsequent papers using *NrlpCre* and *NrlpGFP*. Through identification that activity of the 2.5kb element used in *NrlpGFP* and *NrlpCre* is in fact not limited to rods, it is postulated that previous work using these mouse lines mischaracterized developing cone photoreceptors as rods. In addition, analysis of previously reported datasets also suggests that there is likely transcriptional regulation of *Nrl* in developing cones, and to some extent, protein expression as well. In light of this evidence that the *Nrl* promoter, in both a transgenic and native context, is highly active in developing cones, this suggests that the mechanism by which *Nrl* affects rod/cone fate and/or differentiation is not through differential transcription of *Nrl* and that other mechanisms should be investigated.

## Methods

### Animals

Experiments involving animals were approved and conducted in accordance with animal care protocols of the City College of New York, CUNY. The *NrlpCre* strain houses a transgenic insertion of a 1.7kb fragment of the *Nrl* promoter as well as non-coding exons 1 and 2 driving Cre recombinase (Strain ID: C57BL/6J-Tg(Nrl-Cre)1Smgc/J, Strain #: 028941, RRID: IMSR_JAX:028941). The strain was obtained from Jackson Labs through Charles River (Kingston, NY). *Nrl-Cre* mice were used as heterozygotes. Charles River also provided *Ai14*, CD-1 and C57BL/6J mice. The *Ai14* Cre reporter mice have a loxP-flanked STOP cassette preventing transcription of a CAG promoter-driven red fluorescent protein variant^20^.

### Genotyping

Genotyping was conducted according to protocols specified in the Jackson Labs strain specific Genotyping Protocols document.

### Fixation and Immunohistochemistry

Retinas were fixed for 30 minutes at room temperature in 4% paraformaldehyde in 1× PBS. They were then cryo-protected in 30% sucrose (in 0.5% PBS) overnight at 4 degrees Celsius before being frozen at -80 degrees Celsius. Cryosectioned retinas were permeabilized in 0.3% Triton X-100 for all stages of the staining process. Donkey or Goat serum were used during blocking steps at 10% concentration. Primary antibodies used are in the following table:

| Primary Antibody | Manufacturer | Catalog# | Concentration |
| --- | --- | --- | --- |
| Rabbit anti-Opn1sw | Millipore/Sigma | ZRB1892 | 1:1000 |
| Goat anti-Opn1sw | Santa Cruz | SC-14363 | 1:500 |
| Mouse anti-Cre | Millipore/Sigma | C7988-200UG | 1:200 |
| Rabbit anti-RFP | Rockland | 600-401-379 | 1:250 |
| Chicken anti-mCherry | Aves Labs | MCHERRY 0020 | 1:500 |
| Mouse anti-Rxrg | Santa Cruz | sc-365252 | 1:30 |
| Rabbit anti-Thrb | Proteintech | 21007-1-AP | 1:250 |
| Rabbit anti-Arr3 | Millipore/Sigma | AB15282 | 1:2000 |
| Goat anti-Nrl | R&D | AF2945 | 1:50 |

### Microscopy

Confocal images were obtained using a Zeiss 800 confocal microscope (Oberkochen, Germany). Fiji/ImageJ (National Institutes of Health, Bethesda, MD, USA) was used to analyze images and for manual cell counting. Images were uniformly adjusted for brightness and contrast across singular conditions using Zeiss Zen 3.12.

### Plasmids

The CrxE/px458::GFP and Nrl KO CRISPR guide plasmids were generated and modified as previously described ^21,22^.

### Electroporation

Ex vivo electroporation experiments were performed as previously reported^23^. 8 washes of 70 µL 1× PBS were used to wash the electroporation chamber prior to use and following electroporation of each condition. Electroporation mixes were 50 µL total volume with a 1× PBS final concentration. Plasmid DNA was added to each mix at a 0.2 µg/µL concentration. For the electroporation itself, the electroporator parameters were 25-volt pulses, a pulse length of 50 ms, pulse intervals of 950 ms, and five pulses total.

### Explant Culture

Retinas were cultured on 13-mm/0.2-micron floating filters (10417001; Cytiva [Marlborough, MA, USA]) on Dulbecco’s modified Eagle’s medium/F-12 media (11320082; Gibco [Waltham, MA, USA]) with 10% fetal bovine serum (Thermo Fisher Scientific [Waltham, MA, USA], A3160602) and 1× L-glutamine, penicillin, and streptomycin (10378016; Gibco).

### EdU Pulse and Detection

Postnatal day 0 pups were injected subcutaneously with 20 μL 4 mM EdU resuspended in 1× PBS. Click-iT EdU Alexa Fluor 647 imaging kit (C10340, Invitrogen, Carlsbad, CA, USA) was used to detect the EdU+ cells from CD-1 retina, while EdU+ cells in retina from C57BL/6J mice utilized the following protocol: after immunostaining, slides with cryosectioned retinas were washed three times with 0.3% Triton X-100, subsequently washed once with 1 X PBS, and incubated with an EdU labeling reaction solution for 30 minutes at room temperature. The EdU labeling reaction solution was made by mixing two reaction tubes. The first contained 215 mL 1 X PBS mixed with 10 uL CuSO4 and 0.5uL Alexa647-azide. The second tube contained 22.5uL of RO water mixed with 2.5 uL ascorbic acid at (0.5 M). For each slide, the two tubes were mixed via pipetting and applied for the incubation period before being washed once with DAPI in 0.3% Triton X-100 in PBS (1μg/mL), and a final 1 X PBS wash. This is a modified version of a previously published protocol ^24^.

### Single-Cell Analysis

Single-cell RNA sequencing data was analyzed in **RStudio** using the **Seurat** package. Quality control was performed by filtering cells based on the number of detected genes, transcript counts, mitochondrial transcript content, and predicted doublets. Data was log-normalized and highly variable genes were identified, and then a principal component analysis (PCA) was performed on the scaled data. The number of principal components used for downstream analyses was determined using standard Seurat quality metrics. Unsupervised clustering was performed using a shared nearest-neighbor graph and clusters were visualized using Uniform Manifold Approximation and Projection (UMAP). Differential expression analysis was performed to identify cluster-enriched transcripts. Data visualization was performed using standard Seurat functions, including UMAP, feature plots, violin plots, dot plots, and heatmaps. *Lhx4GFP* RNA-seq data from Buenaventura et al 2019^25^ and the whole-retina RNA-seq from Clark et al 2020^26^ are accessible using GEO Series accession number (GSE132272) and (GSE118614), respectively.

### Analysis of Thrb2Cre scRNA-seq

RNA-seq data from Aramaki et al. 2022^27^ (GSE203481) were retrieved from the NCBI SRA (SRP376331). The FASTQ files were imported into Galaxy.com for further analysis^28,29^. Reads were mapped to the mm10 genome using the RNA STAR tool^30^, incorporating the vM25 GENCODE annotation GTF file to account for splice junctions. The resulting BAM files were visualized using the UCSC Genome Browser. The Salmon quant tool was used to generate tabular files containing transcripts per million (TPM) for each FASTQ file, using an index built from the vM25 GENCODE transcripts FASTA file^31^. The individual cell tabular files were joined to generate a complete expression matrix with genes as rows and individual cells as columns, using the GENCODE vM25 GTF file to annotate the gene abbreviations. These matrices were further analyzed in a Python environment, merging the superior and inferior cells for each timepoint. Briefly, Pandas (v2.3.3) was used for data preprocessing and formatting, and Scanpy (v1.11.5) was used to perform natural log normalization on the TPM values and to generate expression plots^32^.

### Analysis of mouse retina scATAC seq

Previously published scATAC-seq from Lyu et al. 2021^33^ was obtained from GSE181251. The processed scATAC-seq BAM files, aligned to mm10, were downloaded from NCBI SRA (SRP330690) and indexed using SAMtools version 1.23.1. The ‘GSE181251_Single_Cell_ATACseq_cell_annotation.txt.gz’ file provided by the author assigned each cell barcode to its determined cell type. SAMtools view was used to extract reads matching each barcode using the ‘CB’ tag and generate new pseudo-bulk BAM files for each cell type^34^. Cells annotated as ‘early cones’ and ‘cones’ were combined into a single BAM file, as were cells annotated as ‘early rods’ and ‘rods.’ These BAM files were then sorted and indexed using SAMtools. BigWig files were generated using bamCoverage (deepTools version 3.5.6) with a bin size of 50 bp, using a scale factor of (1/cell count) to normalize for variability in cell-type abundance^35^. The resulting BigWig files were uploaded to the Galaxy Europe platform and subsequently visualized using the UCSC Genome Browser.

### Statistical Analysis

Error bars indicate the standard error of the mean (SEM). Statistical analyses were performed using GraphPad Prism 10 (GraphPad Software, Boston, MA, USA) and comparisons between two groups were performed using a two-tailed unpaired Student’s *t*-test with Welch’s correction. *P* < 0.05 was considered statistically significant.

## Results

### The *Nrl* promoter has a history of activity in cone photoreceptors

The *NrlpCre* transgenic mouse has *Cre* recombinase placed under the control of a *Nrl* promoter fragment identical to the one used to make the *NrlpGFP* transgenic. The original publication that characterized this mouse line concluded that *Cre* activity was restricted to rods and not cones although no quantified data was presented in this report^19^. Given this conclusion, we originally obtained this line as a tool that would allow cones and rods to be distinguished during retinal development. Thus, we sought to examine how completely the *NrlpCre* labeled the rod photoreceptor population so as to determine how effectively it could be used to identify rods during development. To do this, we crossed *NrlpCre* positive mice to an *Ai14* loxP reporter mouse line, with the potential to drive TdT fluorescence after recombination **(Figure 1A)**^20^. Adult retinal tissue was harvested from *NrlpCre*/*Ai14* double-positive mice at P60, a stage when the retina is fully mature^36^. Due to the lack of available quantifications of *Cre* activity in rods, and because *Nrl* has been characterized as being rod-specific, we first sought to use the cone-enriched marker Rxrg to mark the cone population so that the Rxrg-negative rod population in the ONL could be quantified^37,38^. *Cre* activity in rods, as determined by TDT+/DAPI+/RXRG-cells in the ONL, showed that approximately ∼90% of rods had current or previous *Cre* activity **(Figure 1B,C)**. This confirmed that the *Nrl* promoter element used in the *NrlpCre* transgenic is widely active in rods, as has been previously published. However, we also observed TdT fluorescence in a number of RXRG+ cone cells. Quantification of these cone cells showed that more than 80% were TDT-positive, revealing they had a history of *Cre* expression **(Figure 1C,E)**. However, in contrast to lineage traced rods, only a small percentage of cones (1.68%) had detectable CRE expression at P60 **(Figure 1E)**. Likewise, identification of cones at P60 using cone arrestin (ARR3+) also showed more than 80% of cone cells were lineage traced **Figure 1D, F)**. A small percentage of ARR3+ cells also had detectable CRE protein (4%) **(Supplemental Figure 1)**. High magnification images of ARR3+ cells confirmed that TDT-cells lacked TDT immunoreactivity in their inner and outer segments. TDT+ cells had TDT immunoreactivity that extended as far as their inner segments, similar to what was seen in rods **(Figure 1G, H)**. Independent of RXRG and ARR3 immunoreactivity, P60 cone nuclei could also be identified by their conventional nuclear organization, which differs from the inverted nuclear architecture of rods^39,40^. Taken together, this suggests that the majority of adult cones do not have current *Nrl* promoter activity and therefore were likely to have activity during earlier stages of differentiation.

**Figure 1.**
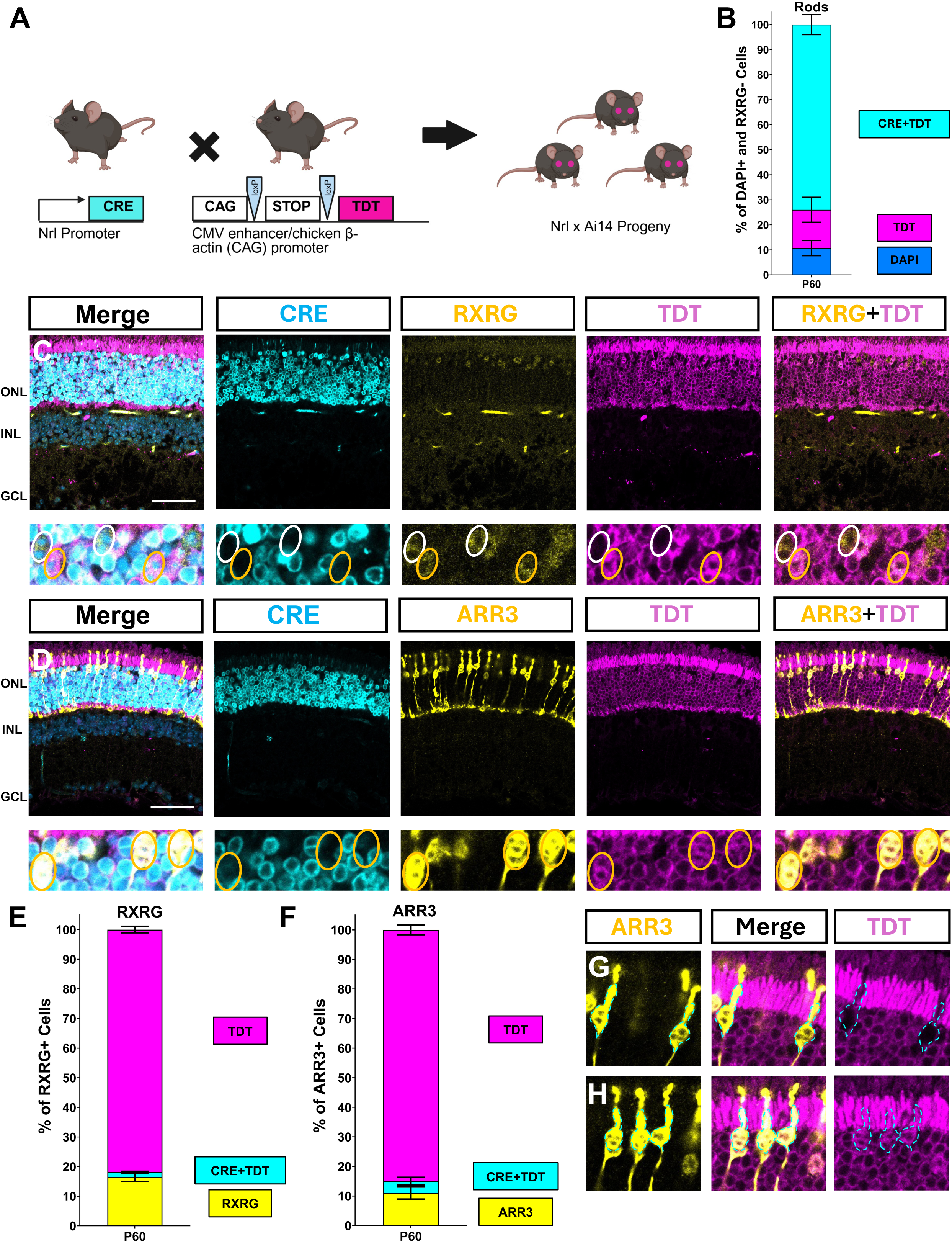
*NrlpCre* lineage tracing labels adult cones as well as rods: (A) Schematic of our experimental workflow. *NrlpCre* mice were crossed with *Ai14* reporter mice and double positive progeny were harvested at time points of interest. (B) Quantification of the percentage of DAPI positive and RXRG-cells in the ONL that are Cre and TDT positive. N=3. (C) Large: single plane images of a P60 *NrlpCre/Ai14* mouse retina stained for CRE and RXRG. TDT signal was amplified using an mCherry antibody. Higher magnification: a single plane with marked cells indicating RXRG+/TDT+ cells (orange circles) and RXRG+/TDT-cells (white circles). (D) Large: single plane images of a P60 *NrlpCre*/*Ai14* mouse retina stained for CRE and ARR3. TDT fluorescence was amplified using an mCherry antibody as in C. Higher magnification: a single plane with marked cells indicating ARR3-+/TDT+ cells (orange circles). (E) Quantification of the percentage of RXRG+ cells in the ONL that express CRE and TDT. N=3. (F) Quantification of the percentage of ARR3+ cells in the ONL that express CRE and TDT, N=4. (G-H) High magnification images of ARR3+/TDT-cells (G) and ARR3+/TDT+ cells (H) from the same image and z-plane shown in (D). Note the dashed lines representing ARR3+/TDT-inner segments in (G) and the ARR3+/TDT+ inner segments in (H). Scale bar in Large images represents 50μm. Higher magnification images are 50μm in length.

### The *Nrl* promoter is active in developing cones

The presence of lineage traced cones in the adult mouse retina, but very few cells actively expressing CRE protein, would indicate that they likely express CRE earlier in their developmental history. Thus, characterization of CRE expression during cone development time points would provide information for the temporal activity of the *Nrl* promoter in cones. To examine this, *NrlpCre*/*Ai14* retinal tissue was harvested at E15.5, during the early phase of cone genesis **(Figure 2A)**, and also at P0, after cone genesis has ceased and they are further along in their differentiation **(Figure 2B)**^41,42^. Quantifications at E15.5 and P0 show that the majority of RXRG+ cells at E15.5 and P0 express CRE protein, with the percentage expressing CRE and TDT increasing between E15.5 and P0 **(Figure 2C)**. We then sought to determine how well the expression of CRE in the *NrlpCre* mouse recapitulated NRL protein. At P1, ∼23% of CRE+ cells were also NRL+ **(Figure 2D).** This suggests that the expression pattern of the CRE protein driven by the *Nrl* promoter is considerably broader than that of endogenous NRL protein. At P1 it was also determined that 100% of all NRL+ cells expressed CRE, validating that any cell that has NRL protein also has *NrlpCre* activity **(Figure 2E)**. This data suggests that during early retinal development the *NrlpCre* transgenic expresses CRE protein within the NRL protein-positive population. However, it also suggests that the activity of the *Nrl* promoter and the expression of CRE protein do not necessitate that a cell will also be NRL protein-positive **(Figure 2F)**. This does not exclude the possibility that CRE may be faithful to *Nrl* on the mRNA level, and there may also be technical limitations restricting our ability to detect NRL protein.

**Figure 2.**
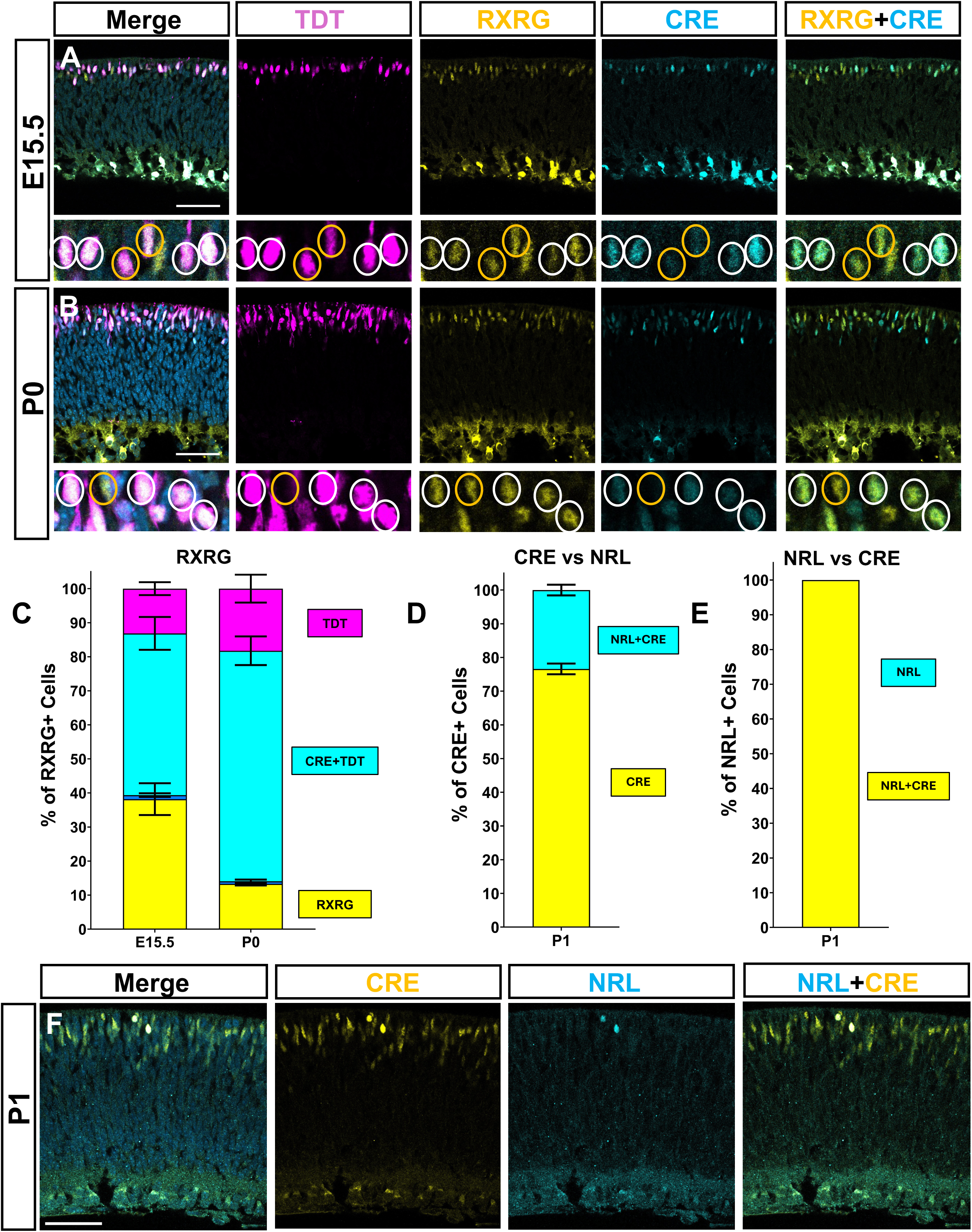
The *Nrl* promoter is active in developing cones at E15.5 and P0: (A) Large: single plane images of a E15.5 *NrlpCre/Ai14* mouse retina stained for CRE and RXRG. TDT fluorescence from the endogenous reporter was amplified using an RFP antibody. Higher magnification: a single plane with marked cells indicating RXRG+/TDT+ cells that are also CRE+ (white circles) and RXRG+/TDT+ cells that appear to have lost CRE expression (orange circles). (B) Large: single plane images of a P0 *NrlpCre/Ai14* mouse retina stained for CRE and RXRG. As before, TDT fluorescence from the endogenous reporter was amplified using an RFP antibody. Higher magnification: a single plane with marked cells indicating RXRG+/TDT+ cells that are CRE+ (white circles) and one RXRG+/TDT- and CRE-cell (orange circles). (C) Quantification of the percentage of RXRG positive cells on the scleral side of the retina that are CRE+ and TDT+ at both E15.5 and P0. N=7 for E15.5. N=4 for P0. (D) Quantification of the percentage of CRE+ cells on the scleral side that express NRL at P1, N=3. (E) Quantification of the percentage of NRL+ cells on the scleral side that express CRE at P1, N=3. (F) Representative images of a P1 *NrlpCre* retina illustrating CRE versus NRL positive cells. Scale bar represents 50μm. Higher magnification images are 50μm in length. Retina orientation: scleral (top), basal (bottom).

We performed CRISPR/Cas9 experiments to disrupt the *Nrl* coding sequence to validate that the *Nrl* antibody could indeed be used as an accurate readout of NRL protein **(Supplemental Figure 2)**. These experiments showed a strong reduction in NRL immunoreactivity in targeted cells, supporting the conclusion that the antibody faithfully recognizes NRL protein in this context.

### Postnatally born rods do not transiently express S-opsin protein

The *Nrl* promoter element has been claimed to have rod-specific transcriptional activity, but the data presented here conflicts with this conclusion and suggests that both rods and cones have *Nrl* promoter activity. This prompted us to reconsider what the impact would be of the *Nrl* promoter’s transient activity in developing cones on studies that interpreted its activity as a dedicated rod readout. One study used the *NrlpGFP* mouse to profile developing *NrlpGFP*+ cells by bulk RNA-Seq and noted that these cells contained cone transcriptional signatures^43^. Under the assumption that *NrlpGFP* is rod specific, this led to the conclusion that rods have a developmental expression of cone genes and this may reflect a transformation of cones into rods as an evolutionary mechanism by which a cone-dominated ancestor gave rise to a rod-dominated descendant. However, if *NrlpGFP* is not rod-specific and is also expressed in cones during development, then the observation of cone genes in the *NrlpGFP* population is simply a result of this population containing rods but also cones. To test whether the Kim et al report correctly identified that a large portion of rods have transient expression of cone genes we conducted an alternative experimental plan to label rods.

This strategy leveraged the fact that in mouse, the vast majority of cones are born prior to birth, while at least half of the rod population is born in the postnatal period^41,42^. Thus, administration of the thymidine analogue 5-ethynyl-2’-deoxyuridine (EdU) to newly born pups can only be incorporated into retinal progenitor cells that generate rods and very few or no cones would be labeled. Thus, it could be determined whether any cell marked by EdU and therefore born in the postnatal period expressed *S-opsin*. To examine this possibility, we injected P0 C57BL/6 mice with EdU to label dividing cells in S-phase and harvested them after 25, 48, and 72 hours **(Figure 3A).** When we stained retinal tissue from these animals for RXRG, S-opsin, and EdU, we found that more than 95% of all S-opsin+ cells at all time points co-stained for *Rxrg*. There were no S-opsin+ cells, out of 3,416 S-opsin+ cells counted, that were co-labeled by EdU **(Figure 3B)**. Thus, at least through the first three days of post-natal development, none of the observed S-opsin+ cells were likely rods born after P0. Additionally, RXRG expression also largely mirrors S-opsin expression, reinforcing the possibility that the S-opsin expressing cells are cones (**Figure 3C-E)**. While Kim et al used a Goat anti-S-opsin antibody (SC-14363), our C57BL/6 quantifications used a Rabbit antibody. A comparison of the immunoreactivities of both antibodies displayed correlative expression **(Supplemental Figure 3)**. As there are differences in S-opsin expression in different mouse strains^44,45^, we also analyzed the Goat anti-S-opsin antibody staining in albino CD-1 mice and obtained very similar results to the C57BL/6 quantifications **(Supplemental Figure 3)**.

**Figure 3.**
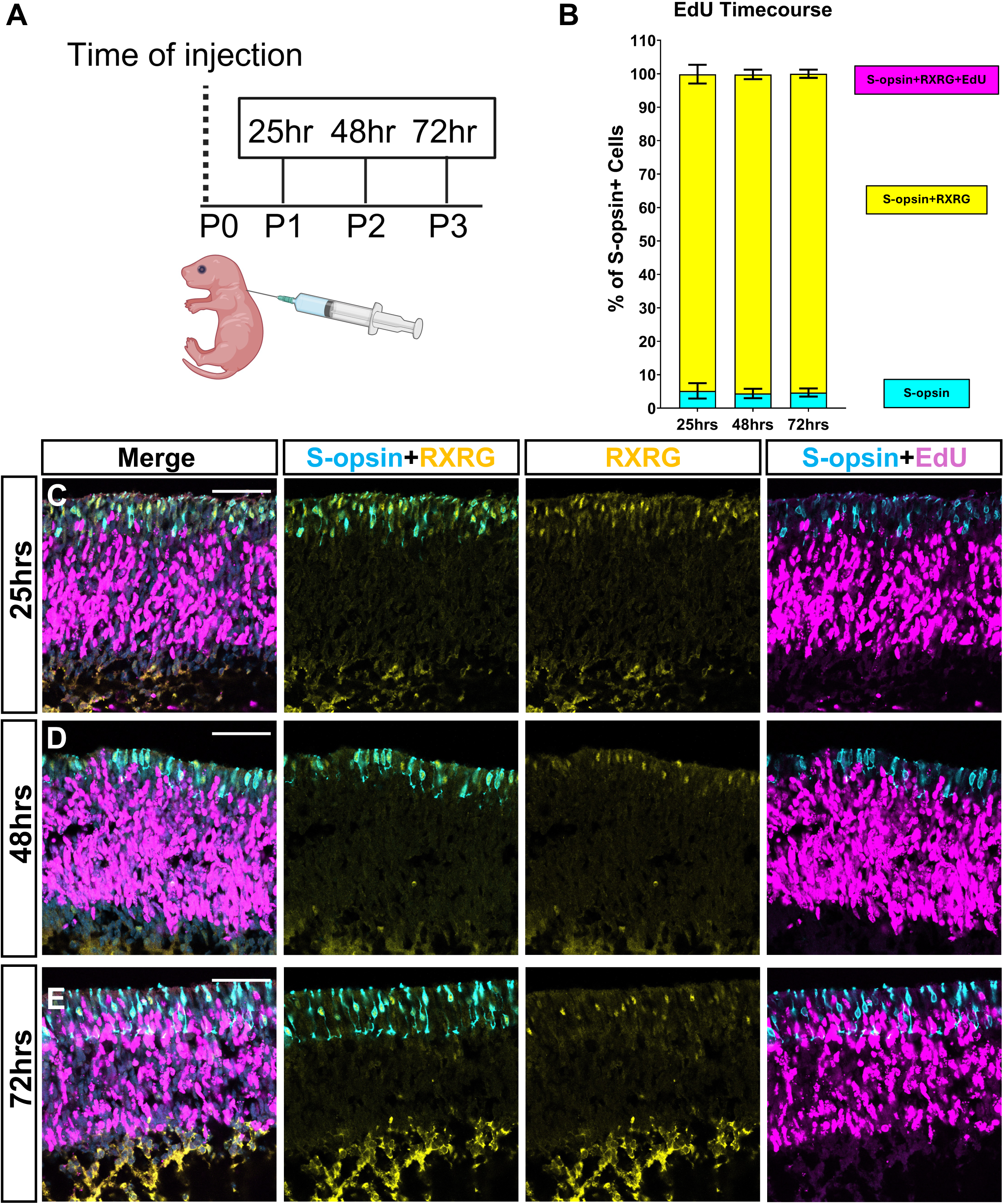
S-opsin positive cells are cones, not P0-born rods: (A) Schematic of the experimental workflow. P0 C57BL/6 mice were injected with thymidine analog EdU (5-ethynyl-2’-deoxyuridine) to label dividing cells. Pups were then harvested at 25hrs, 48hrs, and 72hrs, corresponding with postnatal day 1, 2, and 3, respectively. (B) Quantification of the percentage of S-opsin+ cells that are RXRG+ and EdU+. N=3 for 25hrs, N=8 for 48hrs, and N=6 for 72hrs. (C-E) Representative images of retinas from 25hr, 48hr, and 72hr EdU injected mice. Staining is for EdU, S-opsin, and RXRG. Scale bar represents 50μm. Retina orientation: scleral (top), basal (bottom).

### Endogenous NRL protein is detectable in a subset of developing cones

Since the *Nrl* promoter can drive CRE in developing cone-like cells, and since developing rods born in the early postnatal retina do not seem to undergo a S-opsin expressing phase, we wondered if it was possible that developing cones ever express NRL protein – meaning that the *Nrl* promoter is correctly identifying the presence of NRL protein in cones. A previous report qualitatively and quantitatively reported that in some cells, THRB protein colocalized with *NrlpGFP* during early retinal development^46^. At P0, they also reported occasional *NrlpGFP* and S-opsin co-expression. Thus, we examined P1 retinas of *NrlpCre* mice for expression of NRL, S-opsin, and CRE **(Figure 4A)**. The majority of S-opsin expressing cells at P1 expressed CRE protein, mirroring results seen in Kim et al analyzing S-opsin co- localization with *NrlpGFP*^43^. Interestingly, we found that a number of S-opsin expressing cells also expressed Nrl protein **(Figure 4A)**. *Nrl* and S-opsin co-expression did not co-localize in most cells, with only 10.5% of S-opsin expressing cells possessing NRL immunoreactivity **(Figure 4B)**. To determine whether these S-opsin+/NRL+ cells are rods or cones, the specificity of S-opsin expression in cones was assessed using a published scATAC-seq dataset **(Figure 4C)**^33^. Using the author’s annotations for rod and cone cells, we generated pseudo-bulk clusters for these cell types at various timepoints. Mapping these tracks to the S-opsin locus revealed open chromatin at the S-opsin promoter region at the P0 and P2 timepoints in cones but not in rods. Analysis of a control locus, *Gapdh*, showed similar chromatin peaks between the rod and cone clusters at both time points **(Supplemental Figure 4)**.

**Figure 4.**
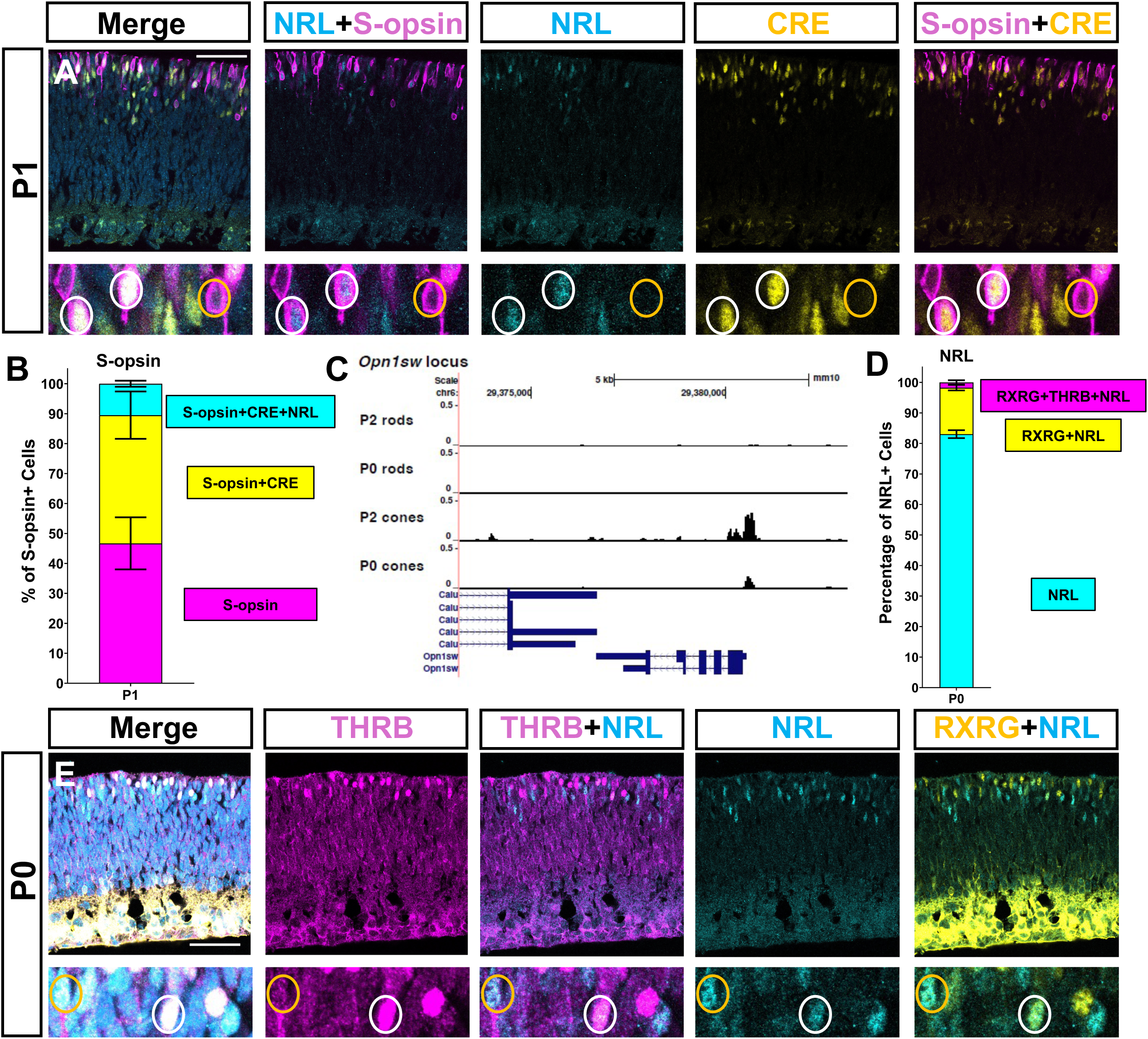
Endogenous NRL protein can be found in cone-like cells during the early post-natal period: (A) Large: single plane images of a P1 *NrlpCre* mouse retina stained for CRE, NRL, and S-opsin. Higher magnification: a single plane with marked cells indicating S-opsin+/NRL+ cells that are also CRE+ (white circles) and an S-opsin+NRL-cell that is also CRE-(orange circle). (B) Quantification of the percentage of S-opsin+ cells that express NRL and CRE in P1 *NrlpCre* mice, N=3. (C) scATAC-seq of P0 and P2 cone and rod clusters with the *Opn1sw* locus in view. (D) Quantification of the percentage of NRL+ cells on the scleral side of the retina that express cone markers RXRG and THRB in P0 NrlpCre mice, N=4. (E) Large: representative images of a P0 *NrlpCre* retina stained for THRB, RXRG, and NRL. Higher magnification: a single plane with marked cells indicating NRL+ cells that are THRB and RXRG+ (white circle) or only RXRG+ (orange circle). Scale bar represents 50μm. Higher magnification images are 50μm in length. Retina orientation: scleral (top), basal (bottom).

NRL co-localization with S-opsin raised the question of whether NRL would also be found in cells expressing other cone markers like RXRG and THRB during the early post-natal period. Quantifications of THRB, RXRG, and NRL expression in P0 retinas showed that a small number of NRL+ cells also co-localized with RXRG+ or THRB+, with 1.72% of NRL+ cells being THRB+/RXRG+ and ∼17% being RXRG+ **(Figure 4D)**. While both *Rxrg* and *Thrb* have been described as pan-cone markers^27,37^, their expression pattern at birth is dynamic, with both genes being downregulated in late-embryonic and early post-natal cones, as has been previously reported^38,47^. Due to our inability to track cones that were in the process of downregulating RXRG and THRB, we limited our confocal imaging to areas that were positive for both cone markers and NRL **(Figure 4E)**. This leaves open the possibility that the true number of NRL+ cones could be under-counted without using a dedicated *Rxrg* or *Thrb* lineage trace.

### NRL protein co-localizes with the cone markers THRB and RXRG at embryonic stages

*Thrb* and *Rxrg* expression is highest during embryonic development when cone genesis is highest^41^. Thus, characterization of whether THRB+ and RXRG+ cells at embryonic time points also express NRL protein will provide some insight into the pattern of *Nrl* expression in cones. Immunohistochemistry of RXRG and THRB with NRL during embryonic time points revealed populations of THRB*+* and RXRG+ cells that co-expressed NRL protein at both E17.5 and E18.5 **(Figure 5A,B)**. For the purpose of quantification, we counted any cell that was THRB+, RXRG+, or THRB+/RXRG+ on the scleral side of the retina as a cone. At E17.5, less than 5% of THRB*+* and RXRG+ cells had NRL co-expression **(Figure 5C)**. This is in line with the Ng et al report characterizing NRL and THRB expression in the developing retina^46^, and also a recent pre-print that found around 4% of Rxrg+ cells in human retinal organoids also expressed NRL protein^48^. At E18.5 the expression patterns of the cone population shifted, and NRL protein was found in just under 18% of the cone population, with RXRG+/NRL+ cells making up the majority of this population at ∼14% of all cones **(Figure 5C)**. Along with the finding in **Figure 4** that some P0 THRB+ and RXRG+ cells express NRL protein, the data confirming *Nrl* expression in embryonic cones suggests the likelihood that some percentage of cones transiently express NRL protein. This is in agreement with the developmental activity in cones of the *Nrl* promoter in the *NrlpCre* transgenic.

**Figure 5.**
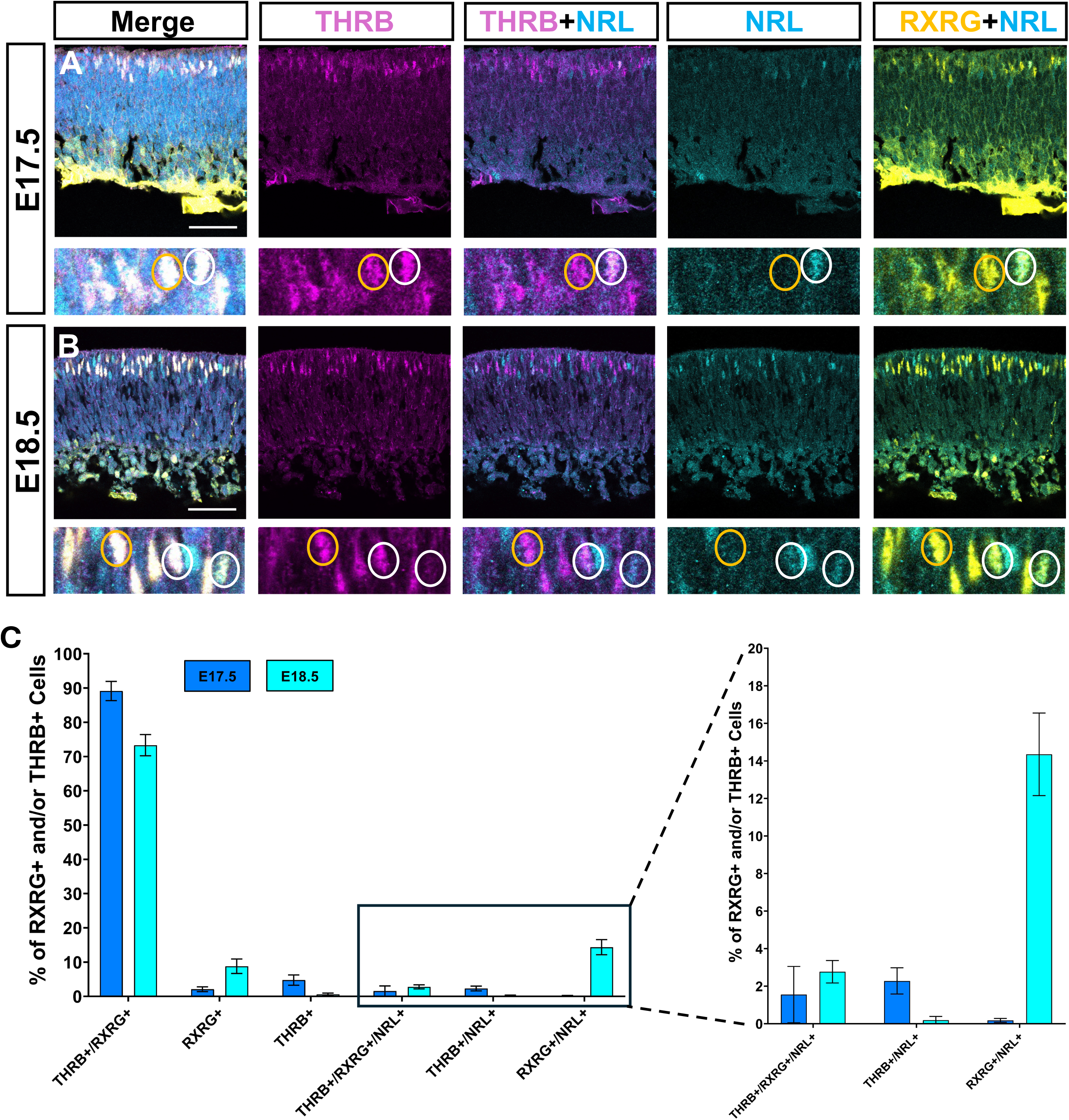
NRL protein co-localizes with cone markers at E17.5 and E18.5: (A) Large: single plane images of a E17.5 CD-1 mouse retina stained for THRB, NRL, and RXRG. Higher magnification: a single plane with marked cells indicating a THRB+/RXRG+ cell that is also NRL+ (white circle) and a THRB+/RXRG+ cell that is NRL- (orange circle). (B) Large: single plane images of a E18.5 CD-1 mouse retina stained for THRB, NRL, and RXRG. Higher magnification: a single plane with marked cells indicating NRL+ cells that are RXRG+ (white circles), one of which has THRB immunoreactivity. A cell that is also NRL- (orange circle) but THRB+ and RXRG+ is indicated (orange circle). (C) Quantifications of the percentage of cells with expression of the cone-enriched genes RXRG and THRB that express Nrl at both E17.5 and E18.5. N=5 for E17.5, N=6 for E18.5. Zoomed graph highlights percentages of THRB+ and RXRG+ cells that express NRL and both time points. Scale bar represents 50μm. Higher magnification images are 50μm in length. Only RXRG+ cells on the scleral side of the retina were counted for both time points. Retina orientation: scleral (top), basal (bottom).

### Explant culture and loss of the RPE elevate NRL expression in cones

Since we also found NRL expression in cone-like cells in vivo, we wondered if cell culture conditions would upregulate NRL expression in mouse retinal explants – recapitulating the upregulation of *Nrl* in cones seen in datasets comparing the chromatin accessibility and transcriptome of the human fetal retina and human retinal organoids^49^. To determine if the behavior of NRL in mouse retinal explants mirrored these changes, we harvested mouse retinas from embryos at approximately E13.5 and cultured them for four days. IHC analysis showed an increase in the number of THRB+ and RXRG+ cells that expressed NRL protein compared to in vivo harvests at E17.5 and E18.5 **(Figure 6A,B)**. In dissected retinas with no RPE, what we have characterized as, “wildtype” cultured retinas, given this is how they are normally cultured within the field, approximately ∼11% of THRB+ and/or RXRG+ cells co-expressed NRL, ∼8% of which were THRB+/RXRG+/NRL+ **(Figure 6A,C)**. The activity of thyroid hormone receptor beta isoform 2 promoter has been recently characterized and lineage traced using a *Thrb2Cre* transgenic in a published report ^27^. This study found that more than 98% of cells with a history of *Thrb2* promoter activity were cones, with only 0.8% of the Thrb2Cre lineage traced population being *NrlpGFP*+. The study characterized those *NrlpGFP*+ cells marked by the Thrb2Cre lineage trace as rods, but there is a possibility that at least some of the Thrb2Cre lineage traced cells that were also NrlpGFP+ are actually cones, given that our own lineage trace data using *NrlpCre* found a small percentage of adult cones still have *Nrl* promoter activity. Thus, because of the specificity of *Thrb2* for cones, it is likely that cells co-expressing *Thrb*, *Rxrg*, and *Nrl* will remain cones.

**Figure 6.**
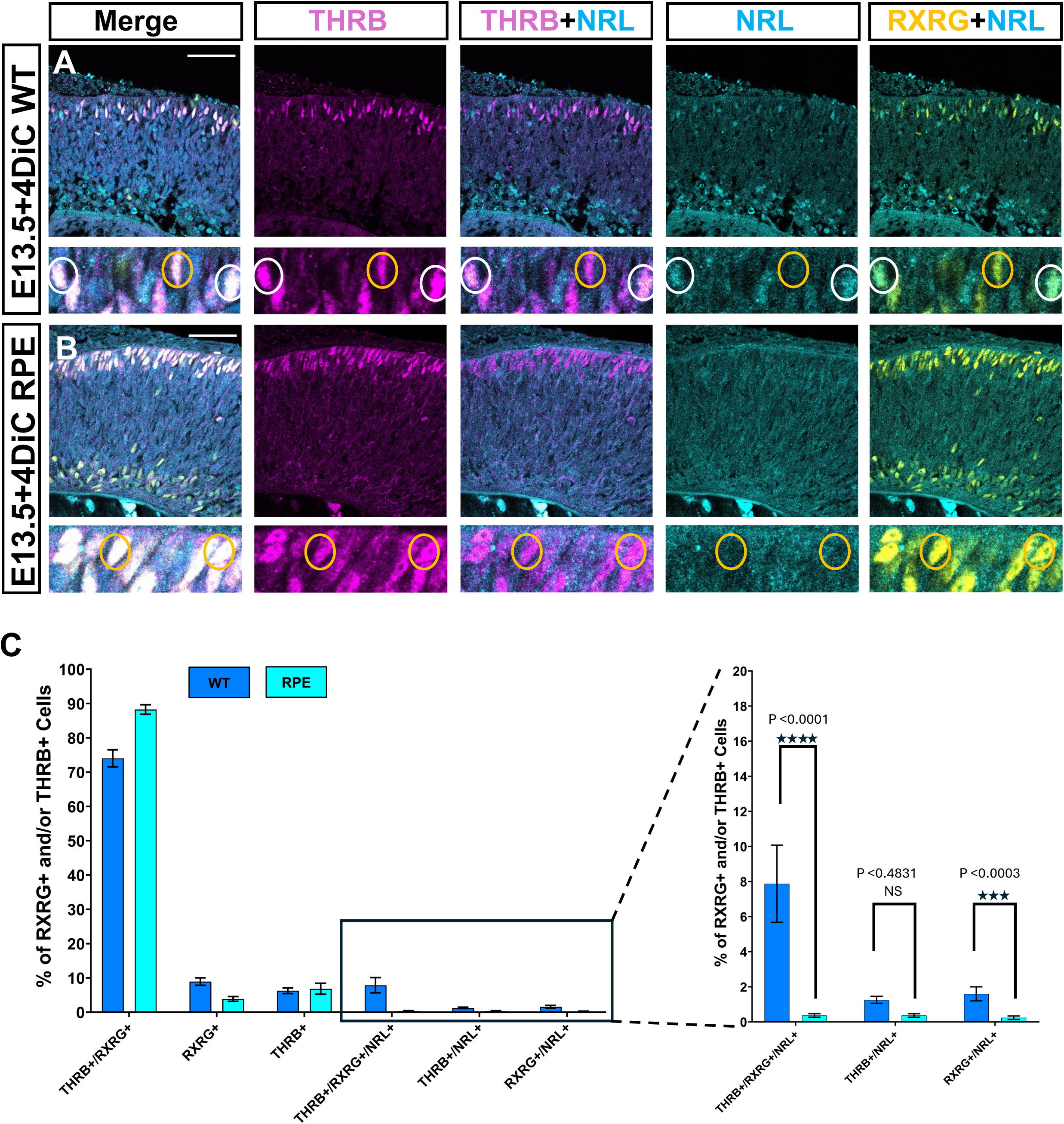
RPE loss elevates cone Nrl expression in retinal explants: (A-B) Retinal explants from E13.5 CD-1 mice were cultured for 4 days (4DiC), with (labeled RPE) or without (labeled WT) RPE. (A) Large: single plane images of a E13.5+4DiC CD-1 retinal explant stained for THRB, NRL, and RXRG. Higher magnification: a single plane with marked cells indicating NRL+ cells that are THRB+ and RXRG+ (white circles) and a THRB+ and RXRG+ cell that is NRL- (orange circle). (B) Large: single plane images of a E13.5+4DiC CD-1 retinal explant stained for THRB, NRL, and RXRG. Higher magnification: a single plane with marked cells indicating THRB+ and RXRG+ cells that are NRL- (orange circles). (C) Quantifications of the percentage of cells with the cone-enriched genes RXRG and THRB that express NRL at E13.5+4DiC without RPE (WT) or with RPE (RPE, N=9). Zoomed graph highlights percentages of THRB+ and RXRG+ cells that express NRL in both conditions. Scale bar represents 50μm. Higher magnification images are 50μm in length. Only RXRG+ cells on the scleral side of the retina were counted in both conditions. Retina orientation: scleral (top), basal (bottom).

The enhanced expression of NRL in culture seems to be a result of altered conditions that do not recapitulate the in vivo environment. One candidate factor we hypothesized may be contributing to the enhanced expression of NRL in culture is the lack of RPE. Thus, retinal explants were harvested at E13.5 and cultured for four days with RPE and surrounding eye-cup tissue intact. After quantifying, it was found that the percentage of THRB*+* and RXRG+ cells that co-expressed NRL dropped significantly, potentially pointing to a role for RPE in regulating NRL expression in in vivo contexts **(Figure 6 B,C).** A disruption to rod genesis was a potential concern, given the lack of NRL expression when retinas were cultured with RPE intact. Thus, retinal explants were also incubated for 6 days in culture to examine rod genesis at later stages of development. At 6 days, NRL expression appeared normally in THRB-/RXRG- cells in retinas cultured with RPE intact **(Supplemental Figure 5).** Similar to 4 days in culture, cells in retinal explants with the RPE removed and cultured for 6 days had frequent co-expression of THRB, RXRG, and NRL **(Supplemental Figure 6).**

### scRNA-seq detects *Nrl* transcripts in *Thrb* and *Rxrg* enriched cone populations

While this report has so far revealed the co-localization of NRL protein with several proteins encoded by cone-enriched genes, we sought to confirm whether *Nrl* RNAs would also be found in cone clusters using scRNA-seq of mouse retinas. A previously published dataset used an *Lhx4GFP* transgenic mouse and flow-sorted GFP+ and GFP- cells from retinas harvested at E14.5 to isolate and enrich for cone photoreceptor precursors for scRNA-seq **(Figure 7A)**^25^. Tissue characterization of GFP+ cells at this timepoint showed a high correlation with RXRG expression suggesting that the vast majority were cone cells. Analysis of this dataset identified that cells with *Nrl* reads were present in the *Lhx4GFP*+/*Lhx4GFP*-combined dataset and cells that were *Lhx4GFP*+ were subset with log-normalized expression values for *Nrl* equal to and above 1 **(Figure 7B)**. The number of *Lhx4GFP*+ cells with *Nrl* represented approximately ∼20% of the total number of *Lhx4GFP*+ cells. As previously reported, *Lhx4GFP*+ cells from this time point are enriched for *Thrb* and *Rxrg* RNAs, as well as many other cone genes **(Supplemental Figure 7)**. Cells that were subset for both *Lhx4GFP* and *Nrl* were also enriched for cone genes, and in particular had high expression of *Thrb* and *Rxrg*, as well as *Gnat2* **(Figure 7C).**

**Figure 7.**
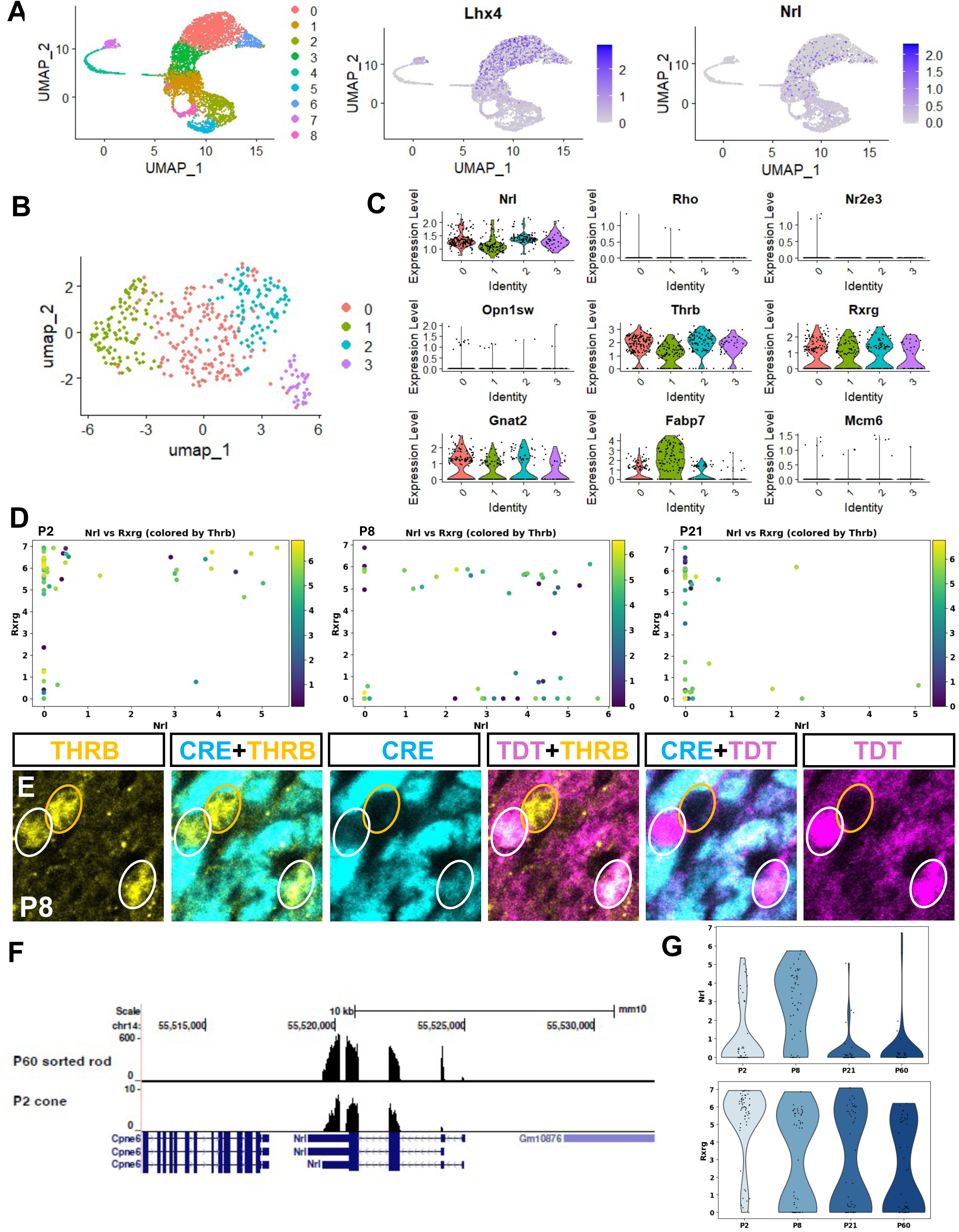
Nrl transcripts in cone-enriched scRNA-seq populations: (A-C) scRNA-seq plots for flow-sorted cells from E14.5 *Lhx4GFP* mice. Cells were sorted such that half were GFP+ and half GFP-. (A) UMAP of all sorted cells present in the dataset. Select feature plots are indicated for *Lhx4* and *Nrl*. (B) UMAP of cells subset for *Lhx4GFP* and *Nrl*. (C) Violin plots of the *Lhx4GFP* and *Nrl* subset for select photoreceptor genes. (D) Scatter plots of hand-picked cells from *Thrb2-Cre*/*Ai6* mice at three distinct developmental time points: P2, P8, and P21. Y-axis indicates *Rxrg* expression while the X-axis indicates *Nrl* expression. *Thrb* expression is illustrated by the color gradient. Units represent the log normalized transcripts per million. (E) Representative single plane images from the retina of a P8 *NrlpCre* mouse highlighting the activity of the *Nrl* promoter element used in both the *NrlGFP* and *NrlpCre* mouse lines. Lineage-traced cells, marked by TDT, that are THRB+ are shown (white circles), with one of these TDT positive cells also retaining CRE expression. A THRB+ cell that does not express either CRE or TDT is also shown (orange circle). (F) Mapped reads at the *Nrl* locus for one of the hand-picked *Thrb2-Cre/Ai6* cells, and a flow-sorted P60 rod from the same dataset. (G) Violin plots of *Nrl* (top) and *Rxrg* (bottom) expression across four time points: P2, P8, P21, and P60. The P60 data was also obtained from Aramaki et al. Imaging is 25μm x 25μm.

To further validate *Nrl* mRNA expression in cones, we utilized a published dataset that isolated *Thrb2Cre* lineage traced cells during the postnatal period to enrich for cones^27^. As the lineage traced cells were reported to be approximately 98% cone-specific, the labeled cells are overwhelmingly enriched for cones. This report showed strong enrichment of cone genes in labeled cells, but *Nrl* was not present in the list of genes with reported reads, though P60 *NrlpGFP*+ cells were also examined. Reanalysis of the raw data found that a population of P2 and P8 lineage traced cells with *Rxrg* and/or *Thrb* reads also had evidence of *Nrl* RNAs **(Figure 7D)**. By P21, the number of lineage traced cones that also expressed *Nrl* largely disappeared. Combined with the E14.5 Lhx4GFP analysis, both datasets potentially point to a developmental progression where *Nrl* RNAs are expressed in early differentiating cones and *Nrl* expression is much weaker or absent as cones mature. To further test the dynamics of *Nrl* transcription and protein expression in postnatal cones, we harvested *NrlpCre*/*Ai14* mice to determine if the *Nrl* promoter indeed remained active at P8. THRB+/CRE+/TDT+ cells were identified confirming the likelihood that *Nrl* RNAs are produced from an active *Nrl* promoter in maturing cones as late as P8 **(Figure 7E)**. While the *Thrb2Cre* dataset we used showed less *Nrl* RNAs at P2 than P8, our P8 immunostaining revealed more THRB and CRE immunoreactivity towards the periphery **(Supplemental Figure 8)**. The authors of the study that produced the *ThrbCre* dataset appear to have been harvesting tissue from areas that were superior and inferior biased **(Supplemental Figure 7)**, likely meaning they harvested retina biased to the periphery. This suggests that P2 peripheral cones likely produce fewer *Nrl* RNAs than cones from the same region at P8.

To confirm that *Nrl* reads were being accurately called in these lineage traced cones, we mapped *Nrl* reads from the P2 *Thrb2Cre* population and compared them to reads from the P60 *NrlpGFP* population. While the P60 *NrlpGFP* sorted rods had much higher *Nrl* reads than the P2 *Thrb2Cre* cones, when we picked out single cells from both populations the *Nrl* reads accurately mapped to the *Nrl* locus **(Figure 7F)**. Violin plots shown indicate *Nrl* and *Rxrg* expression levels in the *Thrb2Cre* dataset at P2, P8, P21, and P60. While *Rxrg* reads remained high in *Thrb2Cre* sorted cells from P2 to P60, *Nrl* is absent in the majority of cells at both P21 and P60 **(Figure 7G)**.

### Whole-retina scRNA-seq and ATAC-seq confirm *Nrl* in the cone lineage

A third published dataset was used to characterize the presence of *Nrl* RNAs in cone-like cells in the context of whole-retina sequencing **(Figure 8A)**^26^. When P2 cells were subset for log-normalized *Nrl* expression equal to and above 1 **(Figure 8B)**, a distinct population of cells that was enriched for cone genes segregated away from most of the other clusters. This cluster, Cluster 4, expressed high amounts of *Opn1sw* and *Gnat2*, as well as some *Thrb* and *Rxrg* **(Figure 8C,D)**. The highest *Nrl* expressing cluster, Cluster 1, had high expression of *Nr2e3*, *Rho*, and *Cngb1*. Cluster 1 had few cone gene reads, likely representing maturing rods **(Supplemental Figure 9).** Feature plots for the *Nrl* subset were produced for *Opn1sw* and *Gnat2* and illustrated that expression of both genes was specific to Cluster 4.

**Figure 8.**
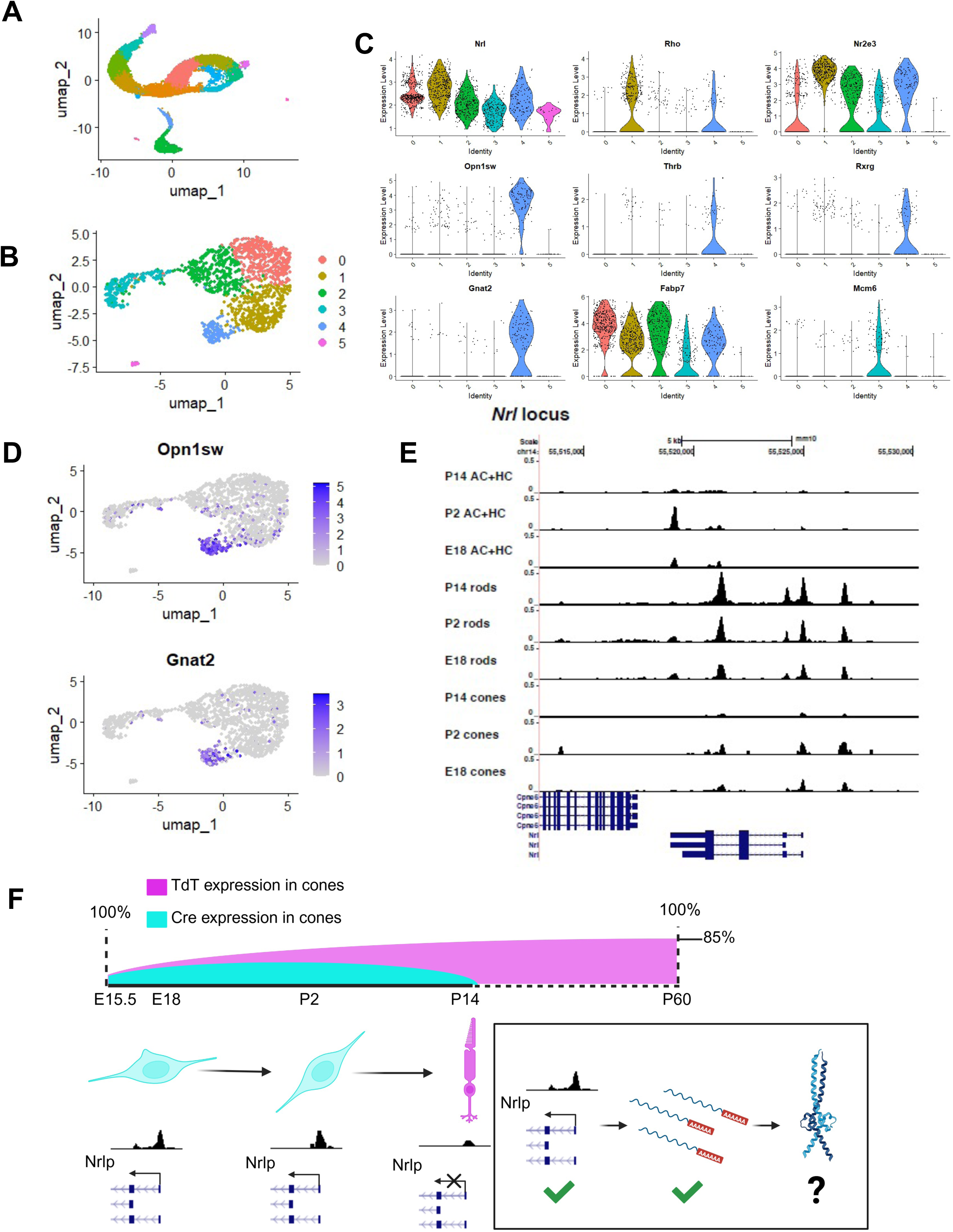
Whole-retina scRNA-seq and *Nrl*-locus accessibility in the cone lineage: (A-D) scRNA-seq plots for cells from a whole-retina P2 dataset. (A) UMAP of all cells from the P2 time point in the dataset. (B) Cells with *Nrl* reads equal to or above 1 were subset from the primary P2 cluster, and a new UMAP was generated. (C) Violin plots of the P2 *Nrl* subset cells for select photoreceptor genes. (D) Feature plots for the cone-enriched genes *Opn1sw* and *Gnat2* are shown. (E) A scATAC-seq dataset was used to assess chromatin accessibility at E18, P2, and P14. Cone, rod, and horizontal/amacrine cell clusters are indicated. (F) Model illustrating the timing of CRE and TDT expression in *NrlpCre/Ai14* mice (top). The timing of CRE expression in the model aligns with *Nrl* chromatin accessibility data at E18, P2, and P14 (bottom left). A simplified model is shown (bottom right): The *Nrl* promoter is accessible during early cone development and *Nrl* RNAs can be produced from the *Nrl* locus during this period. While NRL immunoreactivity was found in cone-like cells, it is unclear how frequently NRL protein is found in these cells and the protein is functional.

RNA contamination can be a significant concern when doing scRNA-seq^50–52^. To reinforce our analysis, scATAC-seq was also considered separately using the same published dataset introduced in **Figure 4**^33^. A comparison of the accessibility of the *Nrl* locus in cones and rods across E18, P2, and P14 demonstrated that E18 and P2 cones have similar *Nrl* accessibility as E18 rods. P2 rods have significantly increased *Nrl* accessibility, and P14 rods have the most **(Figure 8E)**. P14 cones have very little *Nrl* accessibility, in support of our finding in **Figure 7** that maturing cones progressively turn off *Nrl* promoter activity. Accessibility of *Nrl* in horizontal cells and amacrine cells, which have not been reported to express *Nrl* RNAs or protein, is essentially closed at all three time points. The chromatin profile of the control locus *Gapdh* appears to remain relatively consistent across the time points and cell groups considered (Supplemental Figure 10).

## Discussion

Here we report on *Nrl* expression and reporter activity in the developing mouse retina and make three major conclusions: 1) The *Nrl* promoter element in the context of the *NrlpCre* transgenic is active in both differentiating rods and cones, but activity is maintained only in adult rods, 2) *Nrl* transcription occurs in cones during development, but like *Nrl* promoter activity, is also attenuated in more mature cones, and 3) There is some NRL protein expression coincident with cone expressed proteins, in agreement with previous reports, but the fate of these cells has not been confirmed. Below we will discuss the contrast of these results/conclusions with the previous studies that have made conclusions that are relevant to these points.

The Nrl promoter element was reported and characterized in 2006 by Akimoto et al^8^. This study concluded “…that Nrl is indeed the earliest rod lineage-specific reporter.” The conclusion that the promoter element is rod-specific could be drawn from three sources of data in this publication. The primary evidence cited was the adult activity profile of *NrlpGFP* when rods and cones can be clearly identified, and GFP was not observed in cone cells. However, as observed in the current study using *NrlpCre*, *Nrl* promoter activity is detected in developing cones, but seldom in adult ones and so extrapolation across timepoints is not valid. Akimoto et al also included a reference in the text to fate-mapping experiments using the *Nrl* promoter to drive Cre recombinase and stated that their lineage tracing studies supported the finding that the promoter is rod-specific, but no experimental details or actual results were included and data was noted as unpublished. Lastly, introduction of a *Nrl* mutant allele into the background of the *NrlpGFP* reporter showed that *NrlpGFP* cells upregulated cone genes, which supports the idea that *Nrl* represses cone genes and therefore would suggest that *Nrl* expression in cones would be incompatible. However, this genetic data does not necessitate that *Nrl* mRNA expression is specific to rods, as there could be suppression of NRL protein in cones and it is also possible the cone-enriched transcription factor environment may change the gene regulatory networks that *Nrl* participates in. Thus, though Akimoto et al reached the conclusion that the *Nrl* promoter was rod-specific, there was not appropriate rigor applied to support this conclusion and the data that was presented is entirely consistent with the findings presented here: that the *NrlpGFP* transgene has expression in both rods and cones. The subsequent report by Brightman et al generated the *NrlpCre* transgenic line that is used in the current study and their conclusion was that lineage traced cells were specific to rods^19^. We suggest that their conclusion differed from the current study for two possible reasons. The first is that quantitative methods were not used, and so the conclusion was not rigorously supported. Secondly, cone expression of TdTomato was assessed by overlap with cone arrestin staining in the ciliary cone compartment. While we also used cone arrestin staining, we focused our analysis primarily around the nucleus and used nuclear RXRG to look for TdTomato co-localization. In our own microscopy analysis we identified TdTomato expression in the inner segments of cones and rods. We suggest that it is possible the cone highlighted by Brightman et al was one of the approximately 10% of cones that do not have a history of Cre expression, and a more detailed characterization of cone markers versus TdTomato expression markers was needed. Additionally, while they used a Ai9 reporter mouse line, our study used an Ai14 reporter. Thus, it is certainly possible that there could be differences in TdTomato expression between the two mouse lines. However, there is no published evidence to support one of these reporters being more active than the other, and both have been widely used and are considered to be among the best reporter alleles available^20,53^.

The *NrlpGFP* mouse has been used in several studies as a dedicated rod marker, and in cases where this involves a developmental analysis, researchers must reexamine whether cone expression of *NrlpGFP* would impact the conclusions of their study^54–65^. As noted previously, a study where one of the greatest potential impacts of cone expression of NrlpGFP would have on its conclusions, used gene expression profiles of *NrlpGFP* purified cells from developmental timepoints^43^. The observation that *NrlpGFP* cells in the early postnatal mouse retina had mixed cone and rod gene expression signatures while more mature cells did not was taken as evidence that rods transitioned through a cone phase, recapitulating a proposed evolutionary mechanism by which cones were converted to rods to subserve the nocturnal vision needs of mice. This conclusion is predicated on *NrlpGFP* being a dedicated rod marker. However, as summarized above, there is no rigorous data that validates it as such and the current study strongly suggests that the *Nrl* promoter is in fact active in developing cones. The results of the Kim et al study can thus be reinterpreted such that mixed cone and rod gene expression signatures in early postnatal *NrlpGFP*+ cells represents the combined signatures of cone and rod populations that were both labeled by *NrlpGFP* - not a pure rod population. Gene expression profiles obtained at later timepoints would represent pure rod populations. Furthermore, we used EdU birthdating as an independent approach to specifically examine rods, not based on any putative molecular signature, but based on developmental time of origin. This strategy was unable to detect any Sopsin+ cells born in the postnatal period as would be predicted for the rod differentiation pathway proposed by Kim et al. In support of this conclusion, the *Opn1sw* promoter is not in an open chromatin state in developing rod populations, which would be required for transcription. The open chromatin state found at the *Opn1sw* locus in the P2 NrlpGFP flow-sorted cells in the Kim et al study likely represent NrlpGFP+ cones, given that the Nrl promoter is broadly active in cones during the early post-natal period. Lastly, the proposed model of cone conversion to rods is not sufficient on its own to explain the very large population of rod photoreceptors in mice and a diurnal ancestor with a much smaller photoreceptor population, presumably resembling that of extant diurnal vertebrates such as the chicken or ground squirrel. Presumably, other mechanisms that involve increased production of photoreceptors is a much more likely dominant mechanism to account for this evolutionary transition. Taken together, we conclude that the evolution of rod dominance model proposed by Kim et al. is not supported.

Given the findings reported in this study, we propose a model for endogenous *Nrl* mRNA expression in cones whereby the *Nrl* promoter is accessible in early-stage differentiating cones and capable of driving *Nrl* RNAs **(Figure 8F)**. This is supported by the promoter lineage tracing results as well as endogenous scRNA-seq and scATAC-seq analysis. All of the single cell re- analysis included in our study was performed on previously published datasets (including a study from our research group) that did not describe Nrl expression in identified cone clusters. While each of these datasets provides some evidence of this expression, the reproducibility of the phenomenon across them strengthens the evidence. The Aramaki et al dataset is arguably the strongest in this regard because it combines the use of a Thrb2 knock-in Cre transgenic that is validated to be highly cone-specific with deep single cell RNA sequencing^27^. The amount of Nrl transcripts in these cells is much lower than that found in the highly differentiated P60 NrlpGFP rod cells that were also analyzed in this study. However, it is unclear what the comparable level of Nrl mRNA would be in a similarly early-stage differentiating rod because this population was not reported as having been analyzed. The Nrl mRNA expression described here also aligns with a number of recently published reports analyzing scRNA-seq of human fetal retinas and human retinal organoids that reported the presence of Nrl transcripts in cone clusters. A published multiomic dataset comparing accessibility and expression of genes in the human fetal retina and human retinal organoids showed noticeable differences in the accessibility of *Nrl* and the expression of *Nrl* RNAs. Interestingly, the cones analyzed from the human fetal retina had less accessible chromatin at the Nrl locus and fewer Nrl reads than cones analyzed from human retinal organoids^49^.

While we identified cone-like cells that co-express NRL protein, there are caveats to this observation. One is that the THRB antibody used in this study could potentially recognize both characterized isoforms of THRB (THRB1 and THRB2) and only *Thrb2* has been genetically characterized to be cone-specific. However, a previous study that used a THRB2-specific antibody showed co-expression with *NrlpGFP*, which given the proven specificity of *Thrb2* for cones, supports our conclusion that *NrlpGFP* is active in cones^46^. Interestingly, that same study quantitatively found that *NrlpGFP* expression in THRB2 positive cells disappeared by P8, while we found NrlpCre up to P8 but more so in cells at the periphery. Another caveat is that it is still unclear how wide-spread the phenomenon of NRL protein expression in cones is without examination of a dedicated cone-lineage reporter. Detection of NRL protein in cones also does not necessitate that it is functional NRL, which is normally capable of both DNA-binding and transactivation. A NRL protein missing either of those protein domains would be rendered non-functional but could potentially be picked up by an antibody. However, while it is possible the *Nrl* antibody could be picking up a truncated version that lacks transactivating or DNA-binding capabilities, evidence for such an isoform in the mouse retina is not available. It is also possible that *Nrl* expression is not sufficient to repress cone genes in certain cell states. A previous study ectopically expressed NRL in cones and found that in some contexts NRL expression was compatible with S-opsin expression^13^. In addition, no study to date has examined whether NRL is sufficient in cones to repress developmental cone gene expression, leaving open the possibility that it is not. Additionally, the reason for the discrepancy between the numbers of cones with *Nrl* RNA expression and the number of identified cones with NRL protein is not clear. An obvious possibility is that translational regulation could contribute to the dynamic expression of NRL protein and future research would be required to test this hypothesis.

The dominant mechanism proposed for photoreceptor diversification into cones and rods has been centered on *Nrl*. The genetic data is clear that *Nrl* is an essential gene in rod development and in repression of the cone fate and/or differentiation pathway^66–68^. However, the results of this study points to the limits of past studies that have inferred developmental events based on examination of the end result of development. The study of photoreceptor cell fate decisions would benefit from expanded application of approaches that target developmental timeframes and also employ the use of lineage tracing reagents that link developmental events with adult fate outcomes. The results of this study strongly suggest that the mechanism by which *Nrl* exerts specificity in cone and rod programs is not at the transcriptional level. This is important to recognize so that other mechanisms are sought but also because it informs the interpretation of single cell datasets and the use of *Nrl* expression to identify rods. Studies in other model organisms will also be informative to infer more general mechanisms. In the chick, the *MAFA* gene has been suggested to serve as the equivalent Maf factor in rod development as *Nrl* and profiling of developing *THRB*+ cones has failed to identify any *MAFA* expression in these cells^25,69^. Similar results have been observed in zebrafish^43^. Thus, in other species there could be greater transcriptional specificity than in mammals, and the regulatory processes involved in mammalian Nrl expression could be due to the incorporation of this gene into the already established gene regulatory networks of mammalian ancestors. Lastly, while it is known that most vertebrate species have multiple cone types defined at minimum by specific opsin gene expression, there is little known about whether there are unique developmental processes involved in their formation. In mice, there is evidence for an S-cone population defined primarily by synaptic connectivity, true S-cones, which connect to S-cone bipolar cells^45,70,71^. Thus, perhaps one possibility for why only some developing cones expresses NRL protein is related to the origin of specific types of cones or some other undiscovered aspect of mammalian cone formation. Future work is needed to identify the factors that drive Nrl promoter activity in cones during their development. We propose that examining how cones regulate *Nrl* can give insight into the mechanisms governing retinal photoreceptor cell fate decisions and ultimately provide clarity to disease phenotypes stemming from *Nrl* mutations.

