## Supplementary Figures for "Nrl expression and promoter activity in developing cone photoreceptors"

### Supplemental Figure 1: P60 ARR3+ Cones can Co-express CRE

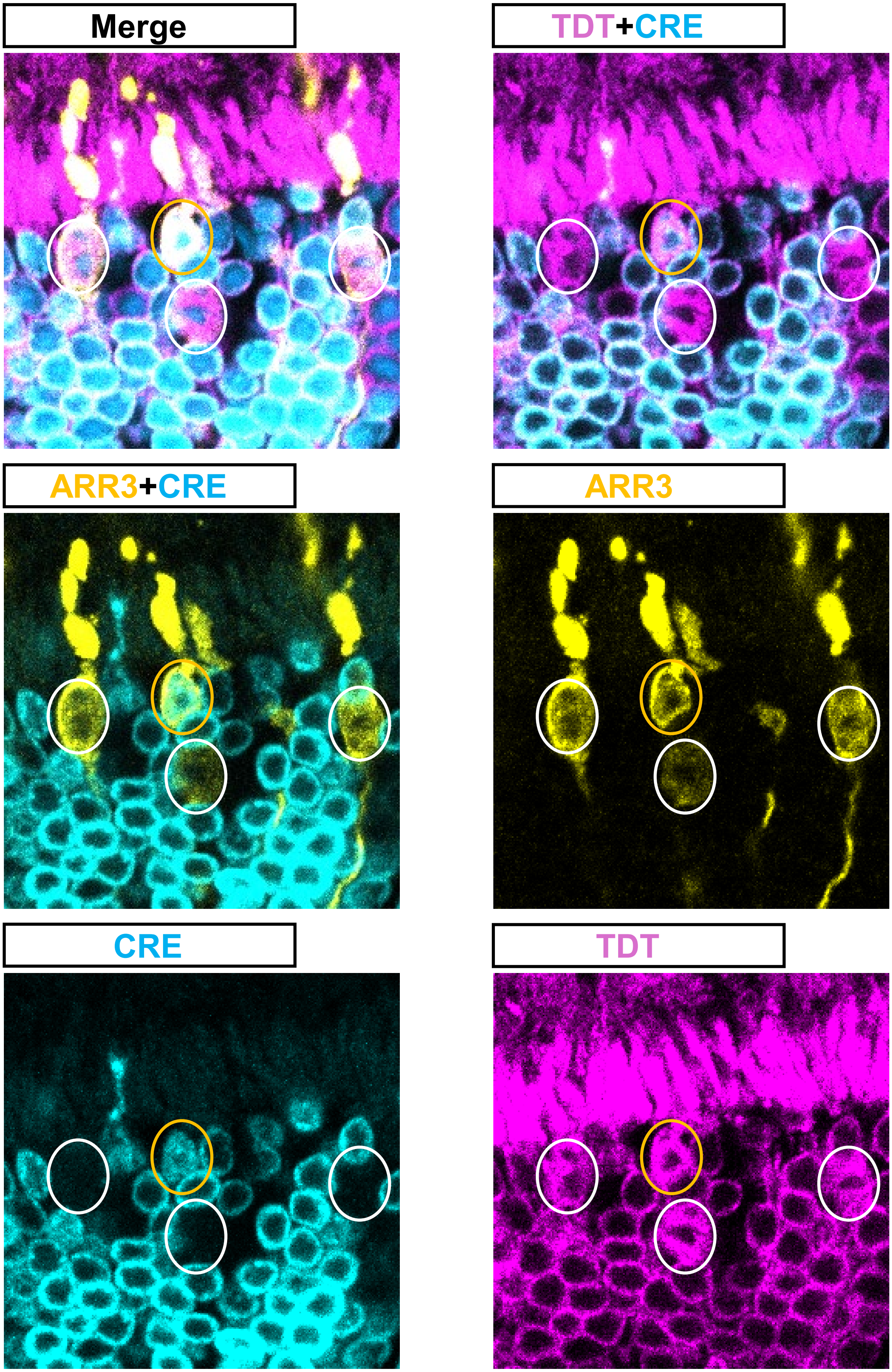

### Supplemental Figure 2:
### Nrl Antibody Verification

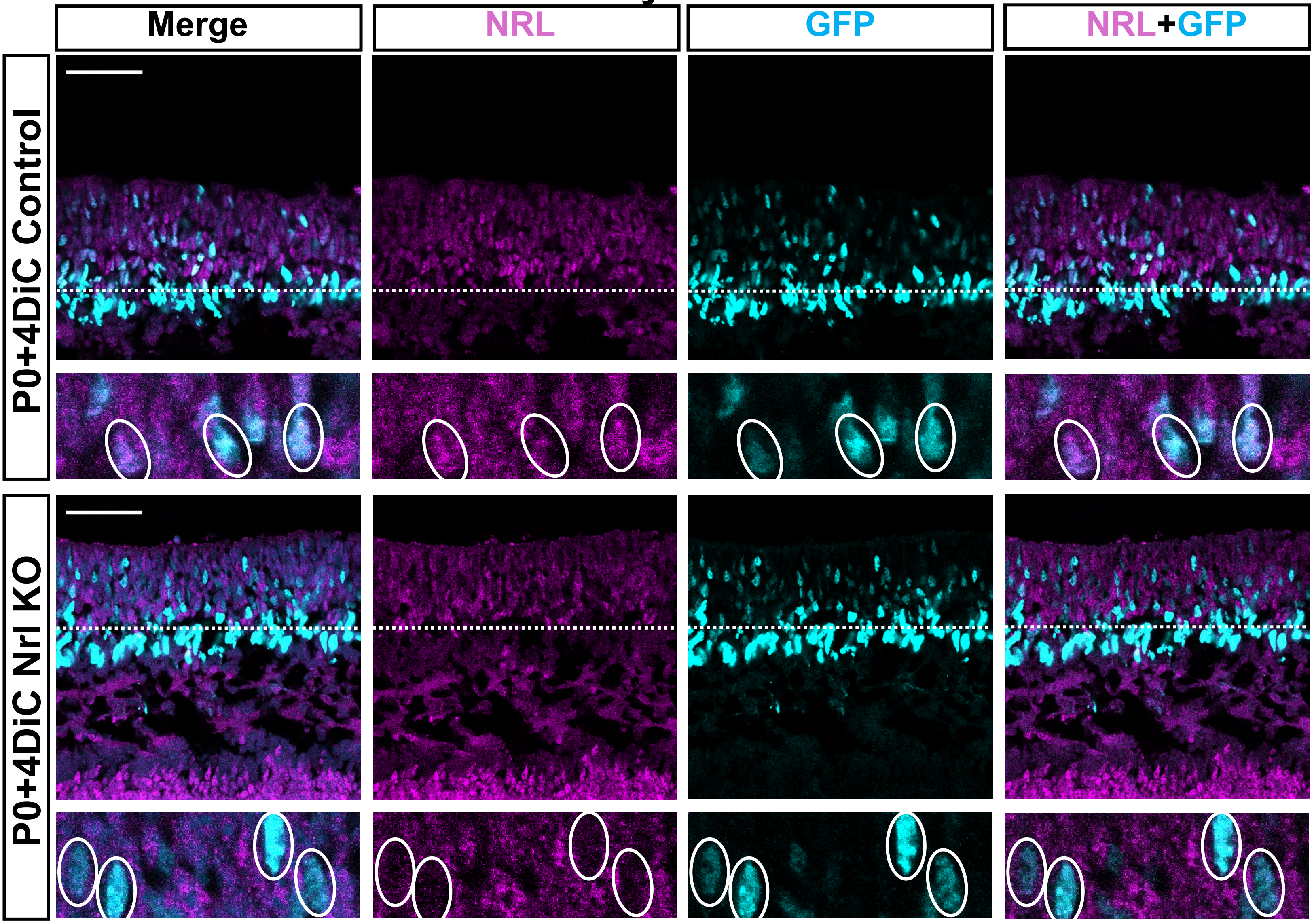


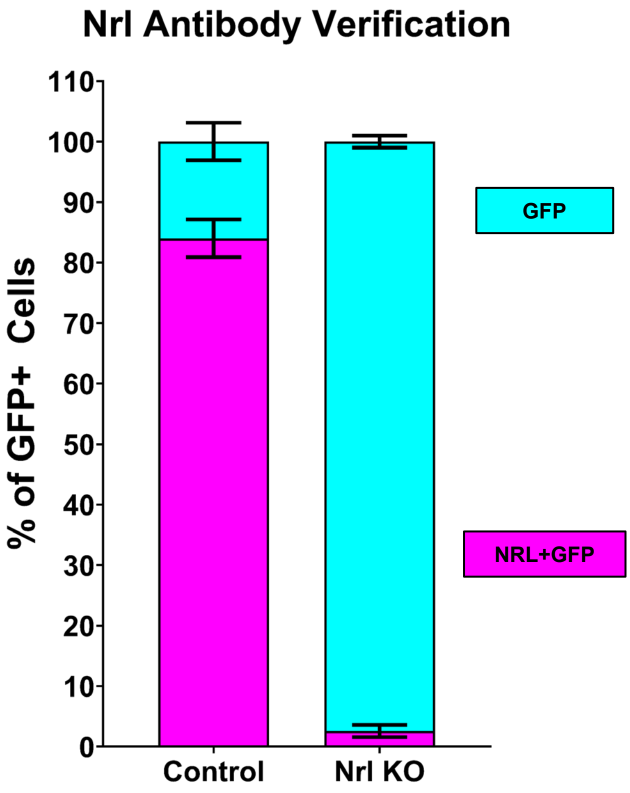

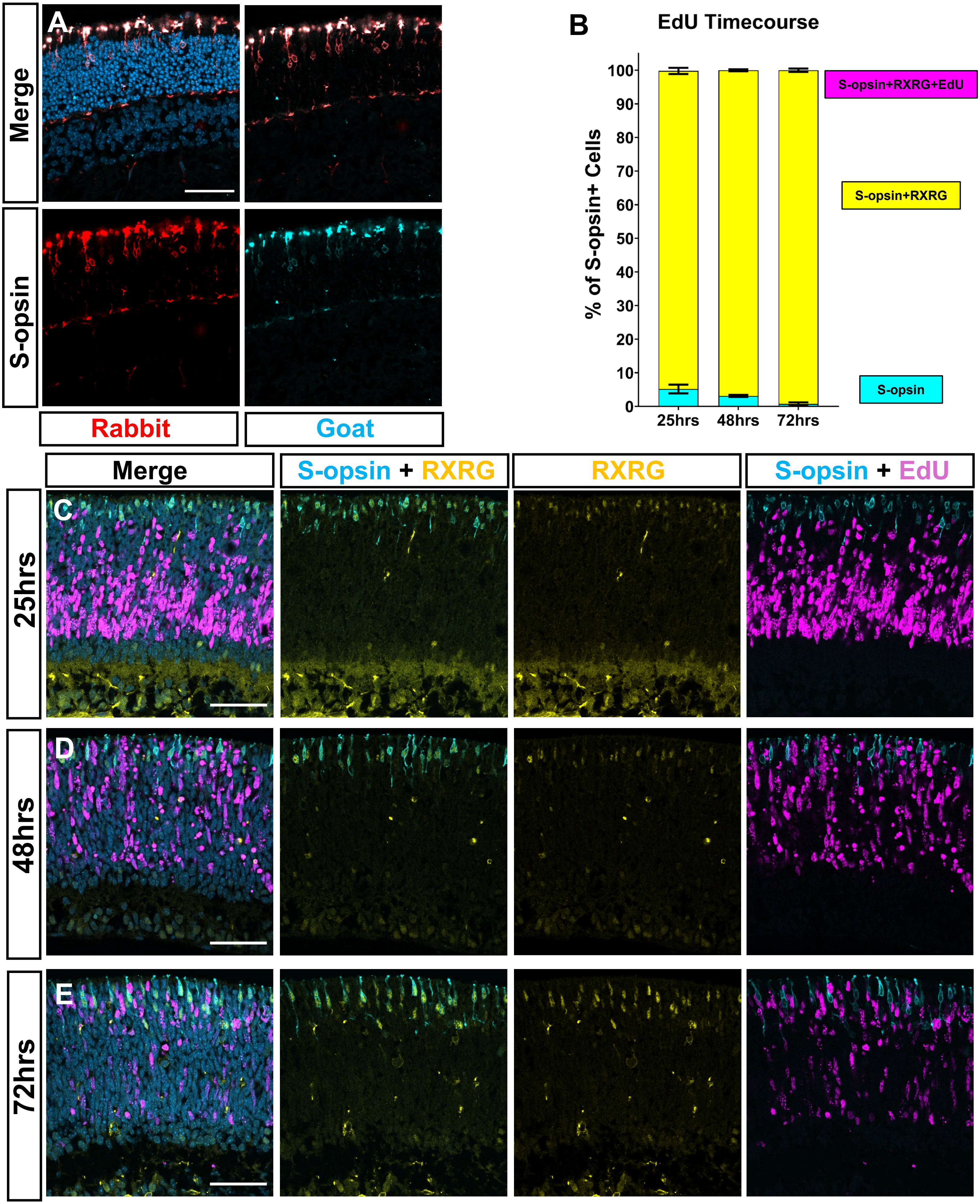
A
Merge
S-opsin
Rabbit
Goat
B
EdU Timecourse
% of S-opsin+ Cells
0
10
20
30
40
50
60
70
80
90
100
25hrs
48hrs
72hrs
S-opsin+RXRG+EdU
S-opsin+RXRG
S-opsin
Merge
S-opsin + RXRG
RXRG
S-opsin + EdU
C
D
E
25hrs
48hrs
72hrs

### Supplemental Figure 4:
### scATAC-seq Control for Opn1sw

*Gapdh* locus

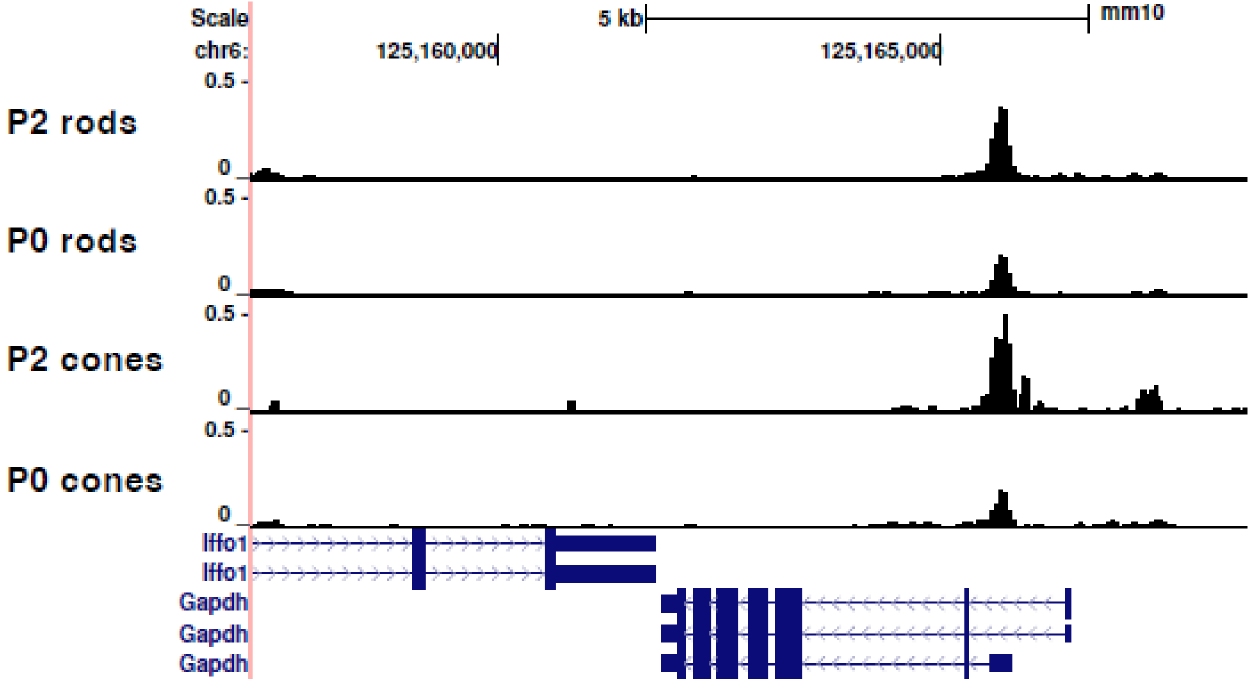

Supplemental Figure 5:

### THRB, RXRG, and NRL Co-localize at 6DiC w/o RPE

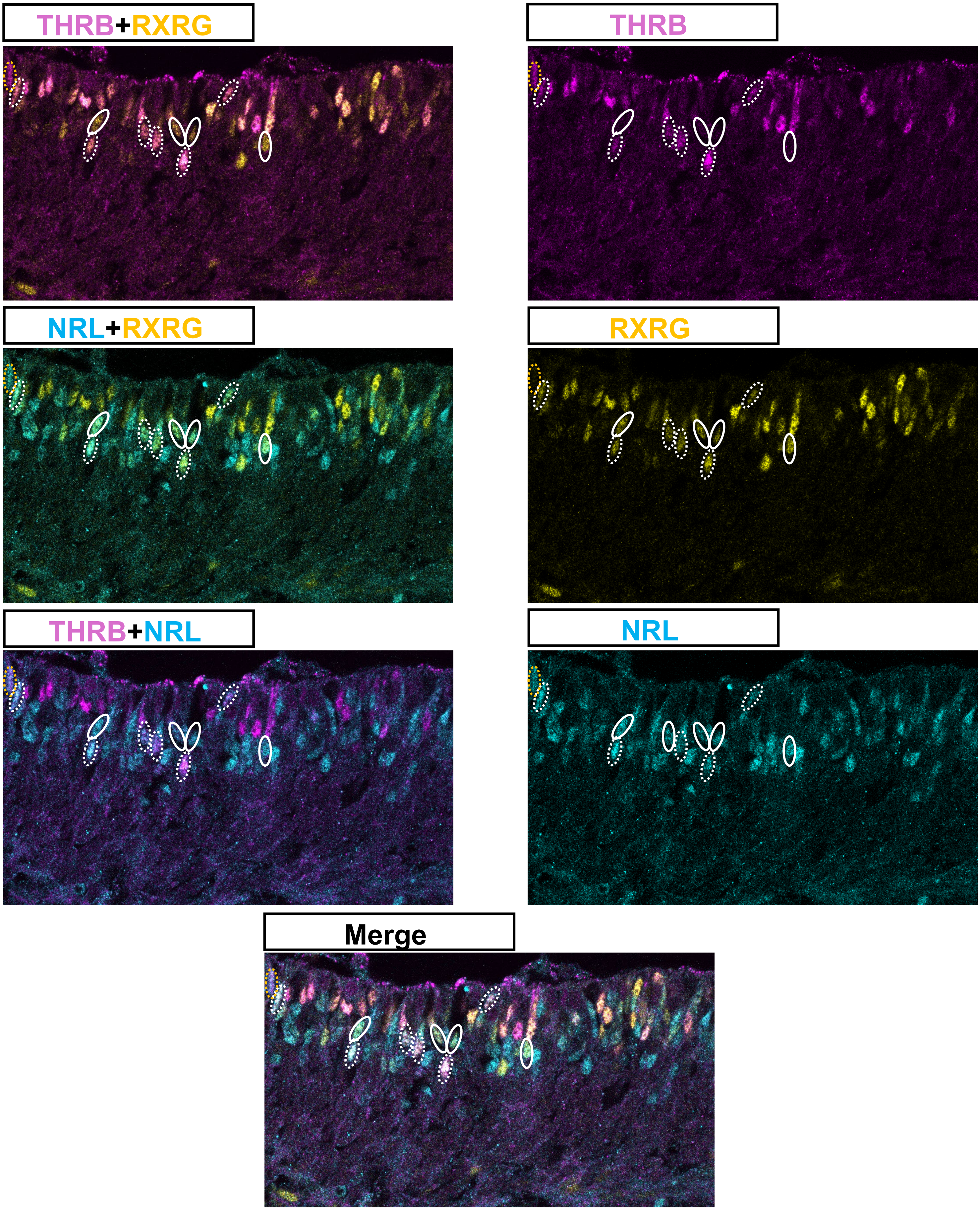

Supplemental Figure 6:
THRB, RXRG, and NRL Rarely Co-localize at 6DiC w/ RPE

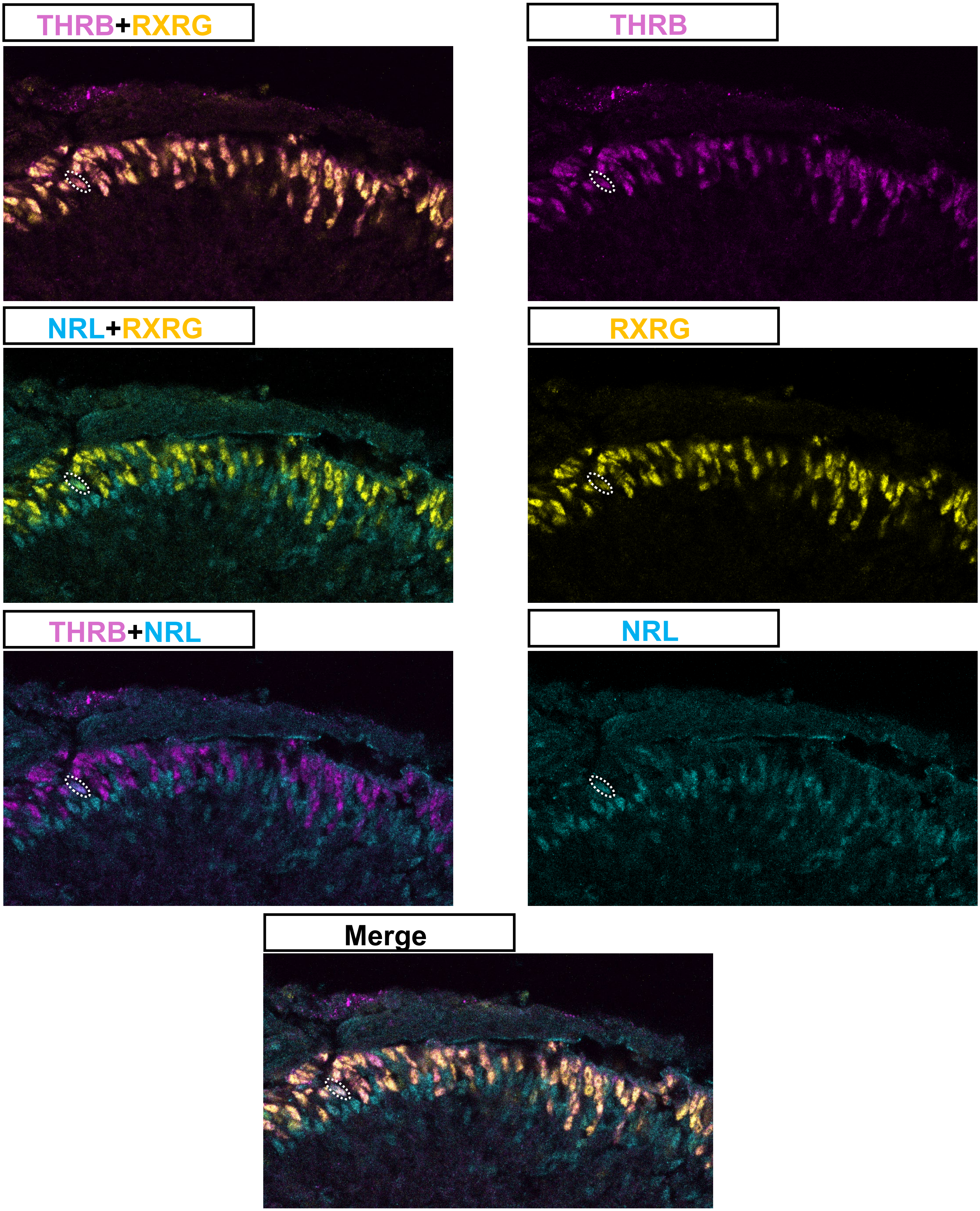

### Supplemental Figure 7: E14.5 Lhx4GFP+

E14.5 Whole-retina

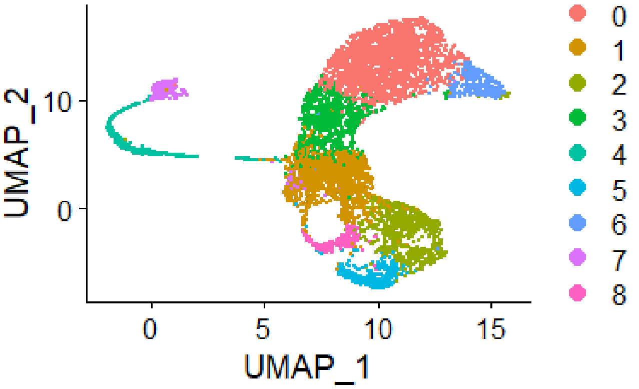


Lhx4GFP+ and Nrl subset

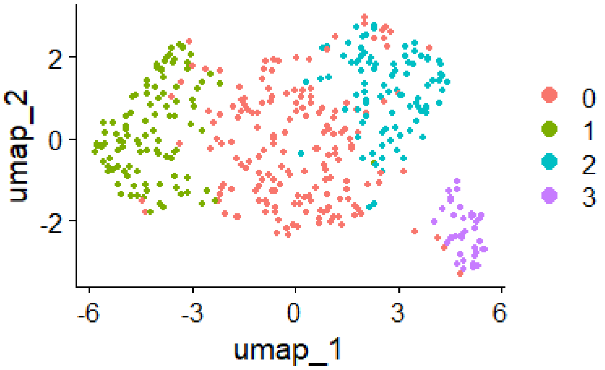


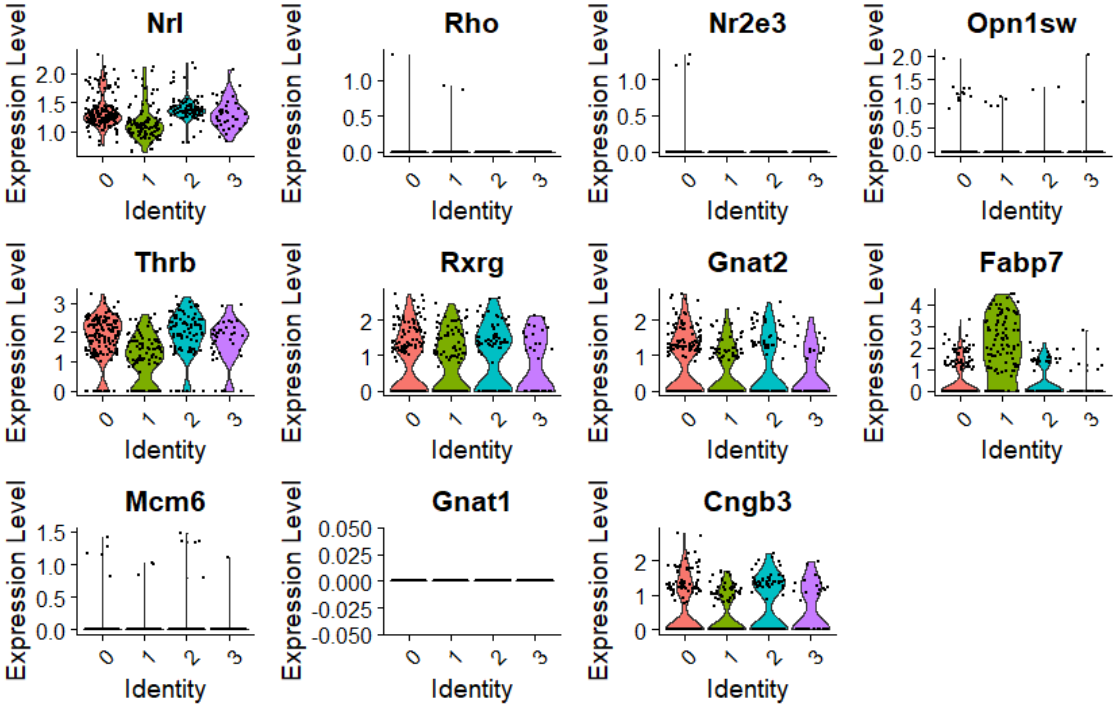


P2 Thrb2Cre

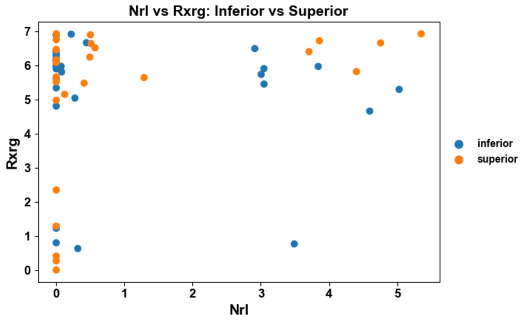


P8 Thrb2Cre

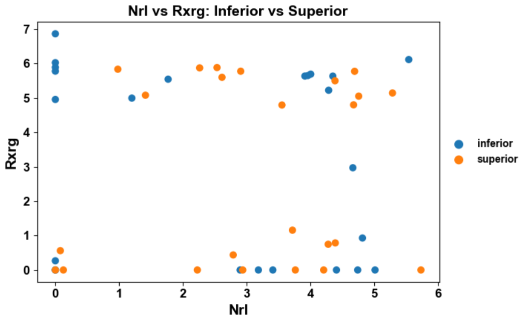


P21 Thrb2Cre

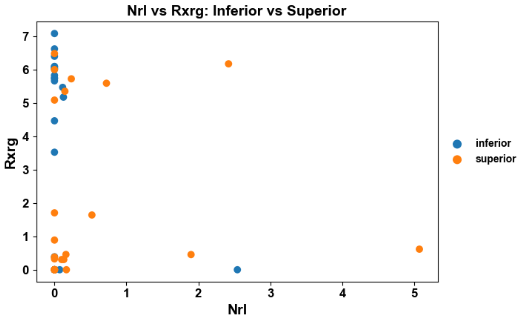

### Supplemental Figure 8:
### P8 THRB+ Peripheral Cones Co-Express CRE

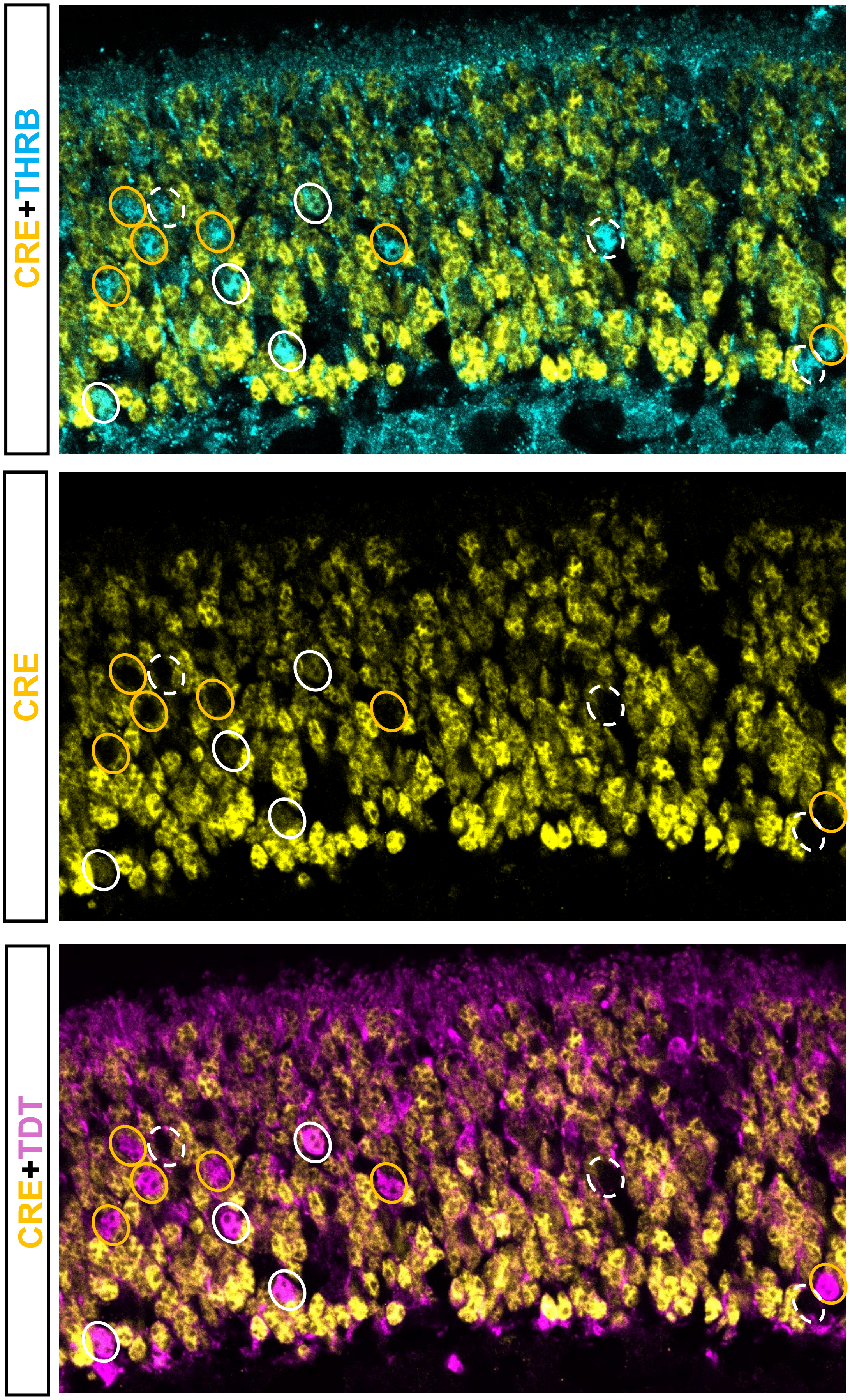

### Supplemental Figure 9: scRNA-seq P2

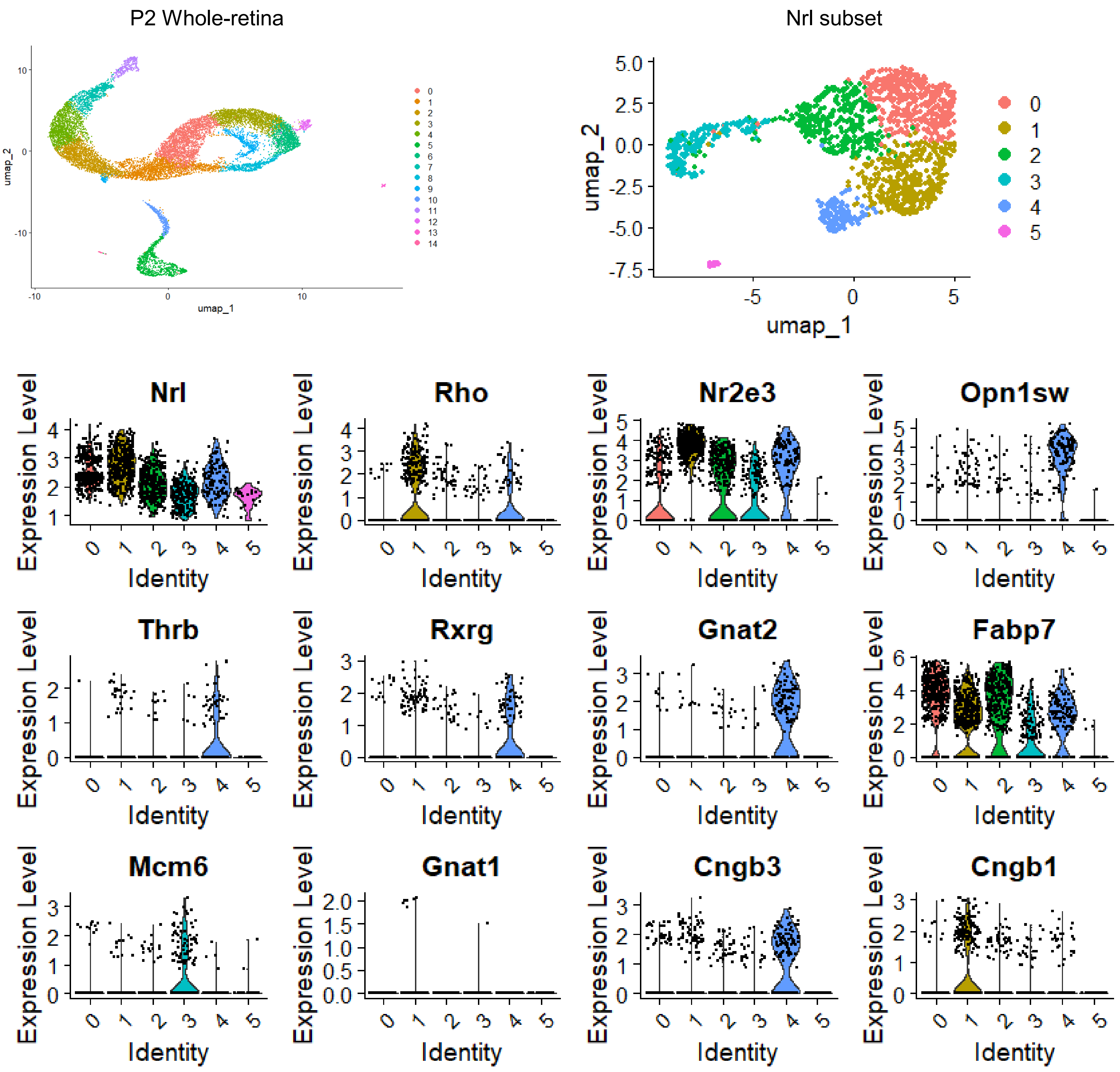

### Supplemental Figure 10: scACAC-seq Control for Nrl

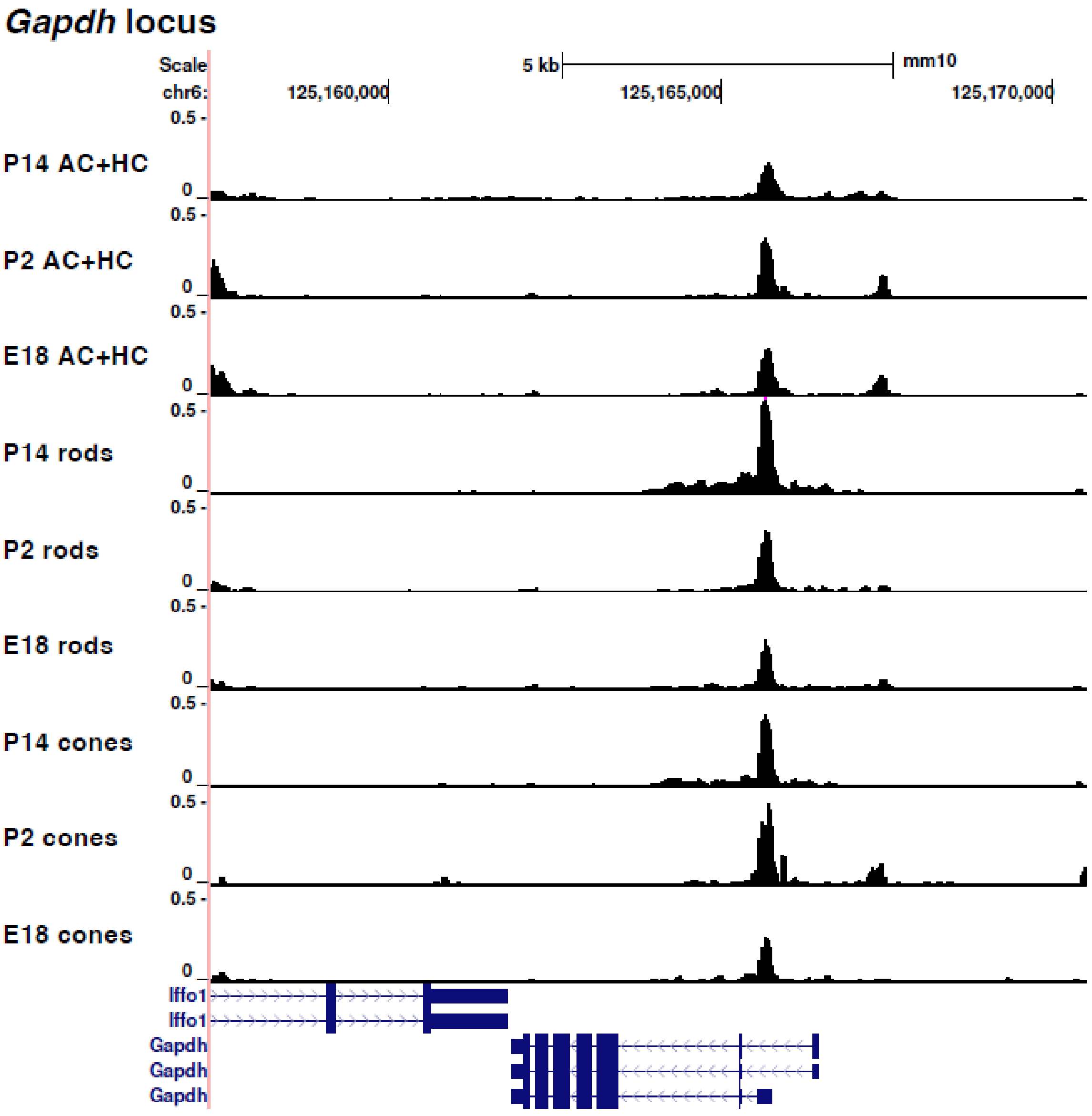

Nrl Manuscript Supplementary Figure Legends

**Supplemental Figure 1 P60 ARR3+ Cones can Co-express CRE:**

Supplemental Figure 1 P60 ARR3+ Cones can Co-express CRE: High magnification images of a representative section of the ONL showing ARR3 and CRE co-localization. The cells also co-express TDT. Orange circles represent ARR3+/CRE+/TDT+ cells. Retina orientation: OS and IS are oriented on the scleral side (top), cell bodies are towards the basal side (bottom). Cropped image is 50μm in width.

**Supplemental Figure 2 NRL antibody verification:**

Supplemental Figure 2: NRL antibody verification: The immunoreactivity of the NRL antibody for NRL protein was tested using a previously characterized Nrl CRISPR KO. It was found that when comparing GFP positive cells between the control and KO vector, the KO vector induced a significant drop in NRL immunoreactivity from the antibody. Scale bar represents 50μm. Higher magnification images are 50μm in length. Retina orientation: scleral (top), basal (bottom).

**Supplemental Figure 3 CD-1 EdU testing:**

Supplemental Figure 3: CD-1 EdU testing: (C-E)Using the CD-1 albino strain, P0 pups were injected and harvested at the same time points as the C57BL/6 experiments. No S-opsin and EdU co-localization was found across the three conditions (B). While the primary figure used a rabbit S-opsin antibody, this staining utilized a goat S-opsin antibody – the same used by Kim et al in their 2016 paper. We checked and found that both antibodies label S-opsin expressing cones 1:1(A). Scale bar represents 50 μm. Higher magnification images are 50 μm in length. Retina orientation: scleral (top), basal (bottom).

**Supplemental Figure 4 scATAC-seq control for *Opn1sw* locus comparison:**

Supplemental Figure 4: scATAC-seq Control for *Opn1sw*: scATAC-seq peaks for *GAPDH* using the same clustering annotations as Figure 4, C.

**Supplemental Figure 5 THRB, RXRG, and NRL Co-localize at 6DiC w/o RPE:**

Supplemental Figure 5: THRB, RXRG, and NRL Co-localize at 6DiC w/o RPE: The degree to which NRL protein was present in cones in culture settings was tested later than 4DiC. At 6DiC, significantly more NRL+ cones were detected. All highlighted cells are NRL positive. Dashed circles represent THRB+/NRL+ cells: the white dashes being RXRG+ and orange being RXRG-. Solid white circles are RXRG+/NRL+ but THRB-. Retina orientation: scleral (top), basal (bottom). Cropped image is 200μm in width.

**Supplemental Figure 6 THRB, RXRG, and NRL Rarely Co-localize at 6DiC w/ RPE:**

Supplemental Figure 5: THRB, RXRG, and NRL Rarely Co-localize at 6DiC w/ RPE: When dissected retinas were culture with RPE for longer than 4 days, there was a significant increase in

NRL protein immunoreactivity. Some THRB+/RXRG+/NRL+ cells were detected, but far fewer than when retinas were cultured in the absence of RPE. A single THRB+/RXRG+/NRL+ cell is highlighted in the representative images. Retina orientation: scleral (top), basal (bottom). Cropped image is 200µm in width.

**Supplemental Figure 7 E14.5 Lhx4GFP scRNA-seq dataset:**

Supplemental Figure 5: E14.5 *Lhx4GFP* scRNA-seq dataset: scRNA-seq plots from the E14.5 *Lhx4GFP* dataset showing a broader number of photoreceptor genes than the original figure. *Cngb1* reads were not found in this dataset, hence its absence. Scatter plots of hand-picked cells from Thrb2-Cre/Ai6 mice are shown at three developmental time points: P2, P8, and P21. Y-axis indicates Rxrg expression while the X-axis indicates Nrl expression. The cells are the same as those in Figure 7D, with identical scaling. Cells collected from the inferior or the superior regions is indicated by blue and orange, respectively.

**Supplemental Figure 8 P8 THRB+ Cones at the Periphery Co-Express Cre:**

Supplemental Figure 6 P8 THRB+ Cones at the Periphery Co-Express CRE: High magnification images of a representative section of the ONL showing P8 THRB+ cells that have evidence of CRE expression. White circles highlight THRB+ cells with CRE+ expression, orange circles are THRB+ cells that are TDT+ but CRE-, and white-dashed circles are THRB+ cells that are neither CRE+ nor TDT+. Images are approximately 115µm in width.

**Supplemental Figure 9 P2 whole-retina dataset:**

scRNA-seq plots from the P2 whole-retina dataset showing a broader number of photoreceptor genes than the original figure.

**Supplemental Figure 10 scATAC-seq control for Nrl locus comparison:**

scATAC-seq peaks for *GAPDH* using the same clustering annotations as Figure 8, E.
